# The BUB1 and BUBR1 paralogs scaffold the kinetochore corona

**DOI:** 10.1101/2025.06.17.660062

**Authors:** Verena Cmentowski, Andrea Musacchio

## Abstract

The kinetochore corona, a polymeric fibrous structure, facilitates chromosome biorientation and mitotic checkpoint signaling during mitosis. How its main building block, the RZZ complex, assembles on the outer kinetochore remains poorly understood. Harnessing corona biochemical reconstitutions and cell biology, we reveal the paralogous checkpoint proteins BUB1 and BUBR1 promote non-redundant branches of corona assembly. MPS1-kinase-dependent kinetochore docking of BUB1 and subsequent recruitment of BUBR1 initiates assembly. Disrupting the first branch by depleting CENP-E, a kinesin that links BUBR1 to RZZ, uncovered a second assembly pathway mediated by a direct interaction between BUB1 and ROD. Discovery of a direct interaction with the RZZ explains how the SAC protein MAD1 fits this corona assembly scheme. Our findings solve the long-standing puzzle of corona assembly and demonstrate the intimate interweaving of chromosome biorientation and checkpoint signaling.

## Main Text

### Introduction

Kinetochores connect chromosomes to spindle microtubules and are essential for chromosome biorientation, mitotic fidelity, and genome stability (*1, 2*). Kinetochores are made of >80 different polypeptides with scaffolding and regulatory functions. The kinetochore corona is a crescent-shaped fibrous structure (*3*). It assembles beyond the KMN (Knl1 complex-Mis12 complex-Ndc80 complex), the main component of the microtubule-binding outer kinetochore layer. Early in mitosis, the corona transports chromosomes as cargoes towards the spindle equator using dynein-dynactin (DD) and CENP-E, microtubule motors of opposite polarity (*4*). Concomitantly, the corona promotes spindle assembly checkpoint (SAC) signaling by docking the MAD1:MAD2 complex (*3, 5*). When Ndc80C dependent end-on microtubule attachments replace motor-based lateral interactions, the corona disassembles, silencing the SAC and licensing sister chromatin separation (*6–8*). CENP-E, the ROD-Zwilch-ZW10 (RZZ) complex, and the DD adaptor Spindly are the corona building blocks. Various outer kinetochore factors partially required for corona stability have been previously identified, but a comprehensive molecular view of corona assembly is missing. Using biochemical reconstitutions, electroporation of recombinant proteins, and separation of function mutants we now describe the entire corona assembly pathway. Our findings reveal that a complex of the paralogous SAC proteins BUB1 and BUBR1 initiates all interactions required for corona assembly. This resolves a long-standing question and underscores why chromosome biorientation and SAC signaling should be viewed as integrated and inseparable phenomena.

### KNL1 scaffolds corona assembly

Assembly of the corona into a fibrous meshwork arises from polymerization of the ROD-Zwilch-ZW10 (RZZ) complex, and also requires binding of Spindly to the RZZ (forming the RZZS complex) and MPS1 kinase-mediated phosphorylation of threonine 13 (T13) and serine 15 (S15) on the ROD subunit (*9–11*) (**Fig. 1A**). We have recently reported that when MPS1 kinase is inhibited with Reversine (*12*) to prevent RZZ polymerization, CENP-E becomes essential for kinetochore recruitment of RZZS (*13*). This, and additional evidence, identifies a previously unrecognized role of CENP-E in corona assembly and disassembly (*14–16*). Electroporation of the 812 kDa recombinant RZZ hexamer (tagged with mCherry) did not rescue corona assembly in cells depleted of CENP-E and treated with Reversine, but electroporation of the equivalent phosphomimetic T13E-S15E (R^EE^ZZ) mutant complex did (**Fig. 1B**) (*13*). Thus, the corona assembles on at least another receptor in addition to CENP-E.

**Fig. 1.**
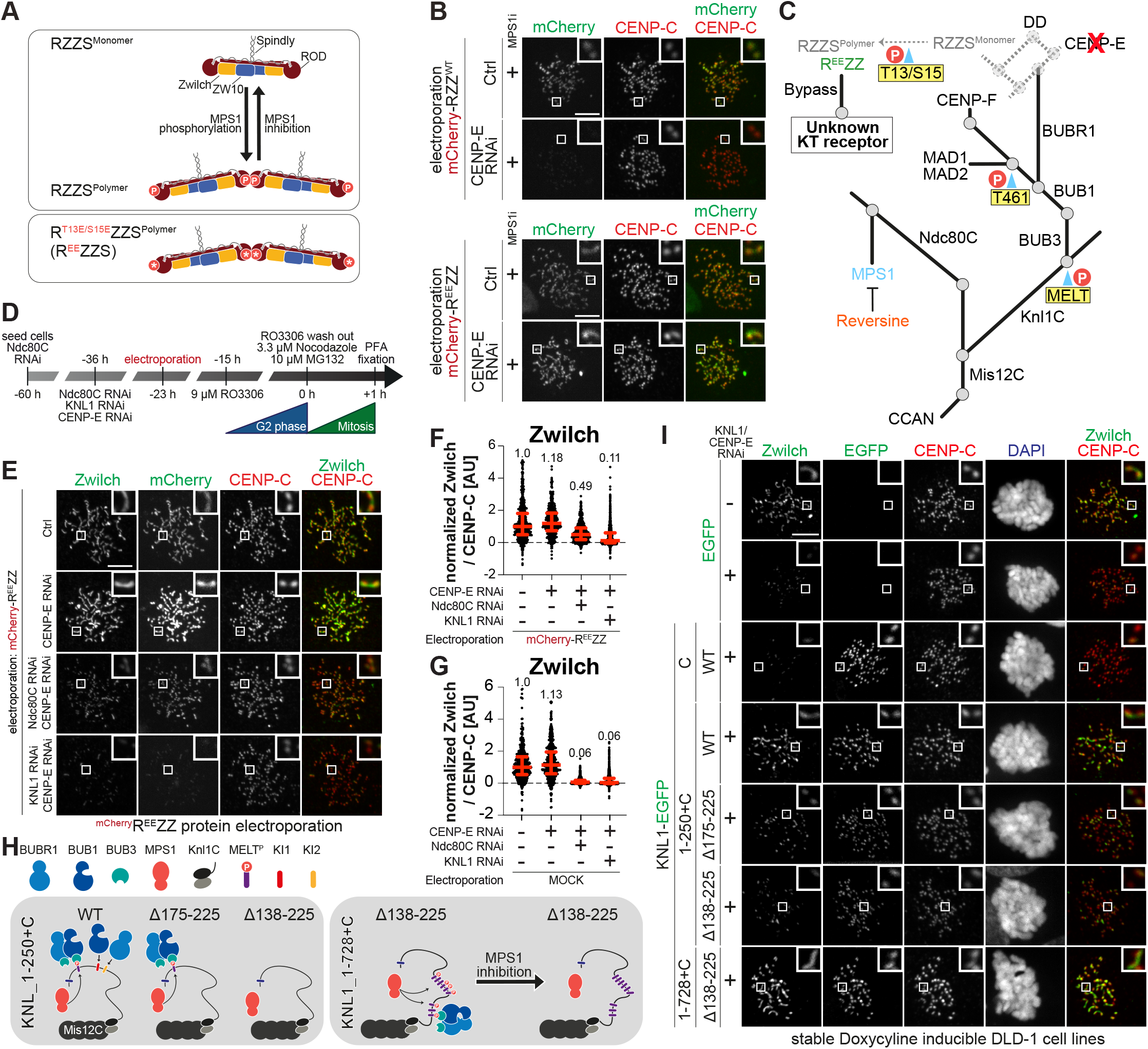
KNL1 promotes corona recruitment. (**A**) Graphical depiction of MPS1-phosphorylation-dependent polymerization of RZZS. (**B**) Representative images showing localization of electroporated mCherry-RZZ^WT^ or mCherry-R^EE^ZZ in cells treated with CENP-E RNAi. Before fixation, cells were synchronized in G2 phase with RO3306 for 15 h and then released into mitosis. Subsequently, cells were immediately treated with 3.3 μM nocodazole, 10 μM MG132, and 500 nM Reversine for an additional hour. CENP-C was used to visualize kinetochores. Scale bar: 5 μm. The experiment was performed three times. (**C**) Organization of outer kinetochore. Each straight line represents a different protein or protein complex. Gray circles represent direct protein interactions. Residues highlighted in yellow are phosphorylated by MPS1. Bypass conditions identified in (B) reveal existence of a second RZZ receptor. (**D**) Scheme for cell synchronization and RNAi treatment for the experiment shown in (E). (**E**) Representative images showing localization of Zwilch in cells electroporated with mCherry-R^EE^ZZ treated for RNAi as indicated and according to the scheme in panel (D). CENP-C was used to visualize kinetochores. Scale bar: 5 μm (**F-G**) Quantification of Zwilch levels at kinetochores of the experiment shown in (E). Red bars represent median (value shown above scatter dot plot) and interquartile range of normalized kinetochore intensity values. n refers to individually measured kinetochores (F left to right: 646, 459, 714, 814, G left to right: 469, 574, 641, 805) from two independent experiments. (**H**) Scheme of the MPS1-dependend recruitment of BUB1/BUB3 and BUBR1/BUB3 to the KNL1 constructs used in the experiment shown in (I). (**I**) Representative images showing the localization of Zwilch in stable DLD-1 cell lines expressing different KNL1 constructs. 24 h after cells were seeded and KNL1 as well as CENP-E were depleted, protein expression was induced through addition of doxycycline and cells were synchronized in G2 phase with RO3306 for 15 h. Subsequently, cells were released into mitosis and immediately treated with 3.3 μM nocodazole, 10 μM MG132 and doxycycline for one hour before fixation. CENP-C was used to visualize kinetochores and DAPI to stain DNA. Scale bar: 5 μm. The experiment was performed three times.

To identify the elusive receptor, we initially assessed how the outer kinetochore affects RZZ recruitment (see schematic in **Fig. 1C**). Simultaneous depletion of the Ndc80 complex (Ndc80C) and of KNL1 by RNAi (directed against multiple subunits in the case of Ndc80C, see Methods) eliminated RZZ entirely, while individual depletions of KNL1 or Ndc80C caused strong depletion, rather than elimination, of kinetochore-bound RZZ (**Fig. S1A-D**). These experiments implicate Ndc80C and KNL1 as being upstream of RZZ recruitment, but do not imply they have a direct role. First, Ndc80C and KNL1 sustain each other reciprocally at the kinetochore (quantified in **Fig. S1C-D**) (*17*). Second, each promotes recruitment of other proteins involved in RZZ recruitment (**Fig. 1C**). Specifically, Ndc80C is essential for kinetochore recruitment of MPS1 kinase (*18–21*), which promotes RZZS oligomerization (**Fig. 1A,C**). Conversely, KNL1 is necessary for recruitment of ZWINT (the second subunit of the Knl1C complex) and BUB1, which is itself MPS1-dependent (*10, 22–30*). Furthermore, BUB1 recruits the MAD1:MAD2 complex (also through MPS1), CENP-F, and BUBR1, whose pseudokinase domain promotes CENP-E recruitment, thereby indirectly contributing to localization of the RZZ complex to the kinetochore (**Fig. 1C**) (*13, 31*). Indeed, depletion of Ndc80C and KNL1 abrogated kinetochore localization of BUB1 and CENP-E in addition to RZZ (**Fig. S1E-F**).

Because mCherry-R^EE^ZZ localizes to kinetochores independently of CENP-E and MPS1 activity, monitoring its requirements for localization might enable us to close in on the RZZ receptor. We electroporated mCherry-R^EE^ZZ into cells depleted of either KNL1 or Ndc80C and monitored its localization with and without MPS1 inhibition (**Fig. 1D**). As CENP-E is not required for mCherry-R^EE^ZZ localization, we took the precaution of co-depleting CENP-E in these experiments to avoid possible confounding effects caused by its role in kinetochore recruitment of endogenous RZZ. Co-depletion of Ndc80C and CENP-E permitted substantial residual mCherry-R^EE^ZZ localization (**Fig. 1E-G**). Further addition of Reversine eliminated residual mCherry-R^EE^ZZ (**Fig. S1G-I)**, indicating that elimination of Ndc80C exacerbates the consequences of MPS1 inhibition by Reversine, in line with the role of Ndc80C in recruiting MPS1 (*20, 21, 32*). As mCherry-R^EE^ZZ bypasses MPS1 in RZZS polymerization, this observation suggests that MPS1 activity is required for an additional aspect of RZZS recruitment. Contrary to the co-depletion of Ndc80C and CENP-E, co-depletion of KNL1 and CENP-E prevented mCherry-R^EE^ZZ recruitment very robustly, and further inhibition of MPS1 had only a minor additional effect (**Fig. 1E-G** and **Fig. S1G-I**).

Collectively, these experiments point to a crucial role of the KNL1 axis in RZZ recruitment, facilitated by MPS1 kinase activity. To identify regions of the 2316-residue KNL1 protein (isoform 2; **Fig. S2A**) relevant to RZZ recruitment, we depleted KNL1 and CENP-E and expressed different KNL1 constructs fused to an EGFP tag in stable DLD-1 cell lines (**Fig. 1H-I** and **Fig. S2C-G**). This analysis revealed that constructs containing at least one of the nineteen MELT (Met-Glu-Leu-Thr) motifs of KNL1 rescued recruitment of RZZ to kinetochores. As MELT motifs bind BUB1:BUB3 upon phosphorylation by MPS1 (*24, 27–31, 33, 34*) (**Fig. 1H** and **Fig. S2B**), this observation argues that BUB1, or a protein downstream of BUB1, contributes to kinetochore recruitment of RZZS in addition to allowing CENP-E recruitment through BUBR1.

To investigate this question, we depleted BUB1 (whose organization is schematically shown in **Fig. 2A**) by RNAi and assessed the impact on RZZ. In comparison to KNL1 depletion, which caused a very strong decrease of kinetochore RZZ, or to simultaneous depletion of CENP-E and BUB1, which completely removed RZZ from kinetochores (quantified in **Fig. 2B**), BUB1 knockdown only caused an incomplete depletion of RZZ from kinetochores (quantified in **Fig. 2B**). These results may seem inconsistent with the known epistatic relationships between the depleted proteins, as depletions of 1) KNL1, 2) BUB1, and 3) BUB1 and CENP-E ought to phenocopy each other with regard to RZZ recruitment (**Fig. 1C**). Nonetheless, previous studies investigating the involvement of BUB1 in RZZ recruitment have also yielded conflicting conclusions (*26, 35–41*), possibly due to incomplete depletions or ectopic interactions (as discussed in Conclusions). Thus, we opted to continue investigating whether BUB1, or of a protein downstream of BUB1, provides a platform for RZZ docking in addition to that provided by the BUBR1-CENP-E axis. Because CENP-E is dispensable for RZZ recruitment when RZZ forms polymers, we limited the analysis to the three proteins known to directly require BUB1 for kinetochore recruitment: CENP-F, BUBR1, and MAD1:MAD2. We have already shown that CENP-F is dispensable for RZZS recruitment (*42*). To test a CENP-E-independent role of BUBR1, we depleted BUB1 and replaced it with a BUB1 transgene lacking the BUBR1-binding region (Δ271-409) (**Fig. 2A**). This mutant continued to recruit RZZS (**Fig. 2C**), thus leaving only MAD1 or BUB1 itself as potential RZZ receptors.

**Fig. 2.**
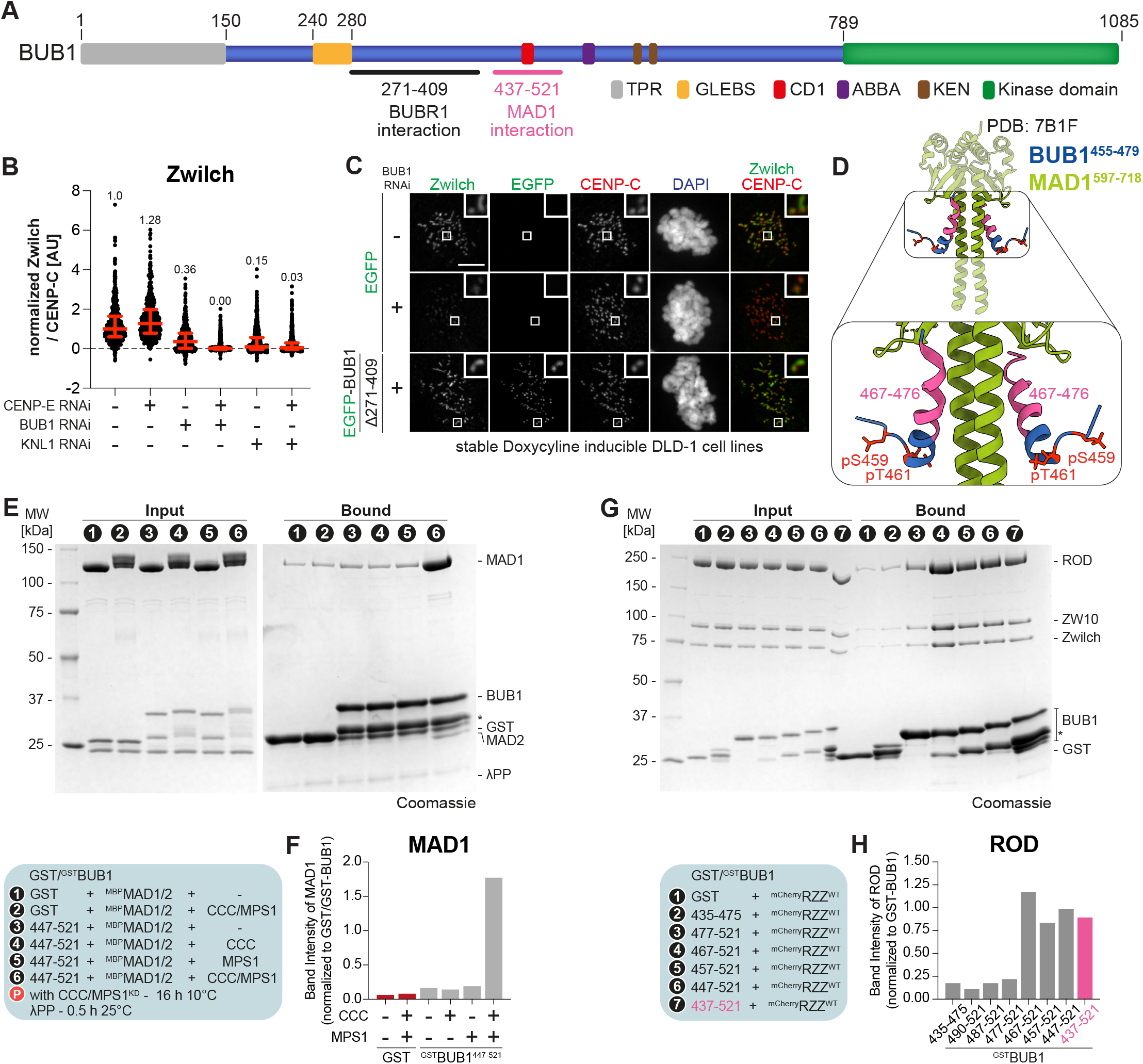
RZZ and MAD1 bind directly to an overlapping region of BUB1. (**A**) Scheme of BUB1 with relevant functional domains. (**B**) Quantification of Zwilch levels at kinetochores of DLD-1 cells depleted either for KNL1, CENP-E or BUB1 as indicated in the figure. 24 h after cells were seeded and treated with siRNA oligos cells were synchronized in G2 phase with 9 μM RO3306 for 15 h. Subsequently, cells were released into mitosis and immediately treated with 3.3 μM nocodazole, 10 μM MG132 for one hour before fixation. Red bars represent median (value shown above scatter dot plot) and interquartile range of normalized kinetochore intensity values. n refers to individually measured kinetochores (left to right: 662, 541, 748, 916, 803, 778) from two independent experiments. (**C**) Representative images showing the localization of Zwilch in stable DLD-1 cell lines expressing different BUB1 constructs. 24 h after cells were seeded and BUB1 as well as CENP-E were depleted, protein expression was induced through addition of doxycycline and cells were synchronized in G2 phase with RO3306 for 15 h. Subsequently, cells were released into mitosis and immediately treated with 3.3 μM nocodazole, 10 μM MG132 and doxycycline for one hour before fixation. CENP-C was used to visualize kinetochores and DAPI to stain DNA. Scale bar: 5 μm. The experiment was performed three times. (**D**) Crystal structure of BUB1^455-479^ and MAD1^597-718^ (adapted from Fischer et al., 2021; PDB: 7B1F). CDK1 and MPS1 phosphorylation sites are highlighted in red. (**E**) SDS-PAGE analysis of a pulldown assay with either GST or GST-tagged BUB1 constructs as bait and MBP-MAD1/2 as prey. For overnight phosphorylation bait and prey were incubated with CCC and MPS1^KD^ for 16 h at 10°C. Before SDS-PAGE analysis the eluate was dephosphorylated with λ-phosphatase for 30 mins at 25°C. (**F**) Quantification of the MAD1 band intensity normalized to the bait signal from the SDS-PAGE depicted in (E). The experiment was performed twice. (**G**) SDS-PAGE analysis of a pulldown assay with either GST or GST-tagged BUB1 constructs as bait, and mCherry-RZZ as prey. (**H**) Quantification of ROD band intensity normalized to the bait signal from the SDS-PAGE depicted in (G).

### BUB1 binds directly to RZZ

The C-terminal domain of MAD1 (MAD1^CTD^, residues 585-718) binds directly to residues 455-479 of BUB1 after phosphorylation of residues Ser459 and Thr461 by CDK1 and MPS1, respectively (**Fig. 2D**) (*43, 44*). Deletion of residues 437-521 of BUB1 encompassing this MAD1 binding site was also shown to reduce RZZ kinetochore levels (*36*). Previous evidence, discussed below, suggests that RZZ acts upstream of MAD1:MAD2 recruitment. Based on our new observations, we hypothesized that BUB1, by binding MAD1, may also facilitate RZZ recruitment. To test this, we first asked if we could reconstitute binding of MAD1:MAD2 to an immobilized GST-tagged fragment of BUB1 encompassing the previously identified binding region (BUB1^447-521^). Indeed, both full-length MAD1:MAD2 and the MAD1^CTD^ bound robustly to this fragment of BUB1 after *in vitro* phosphorylation with MPS1 and CDK1, and preventing phosphorylation abrogated binding (**Fig. 2E-F, Fig. S3A-D**, and **Fig. S5A-B**), in line with a previous report (*43*). In an unanticipated twist, a similar experiment using the same BUB1 bait and RZZ as the prey also revealed robust RZZ binding (**Fig. 2G-H** and **Fig. S3B-C**). We therefore reasoned that MAD1 and RZZ may bind concomitantly to different parts of the same BUB1 fragment. To test this, we titrated increasing amounts of MAD1^CTD^ into a GST-BUB1 binding assay where RZZ was kept at a constant concentration. Contrary to our expectation, the amount of RZZ bound to BUB1 progressively decreased as MAD1^CTD^ concentration increased (**Fig. S3B-D**). Thus, the MAD1^CTD^ and RZZ bind competitively to an at least partly overlapping region of BUB1.

To clarify the molecular basis of this unanticipated competition, we dissected the BUB1:RZZ interaction in more detail. Within BUB1^437-521^, an N-terminal α-helix between residues 465 and 475, coinciding with the ubiquitously conserved central domain 1 (CD1, also called CM1; residues 458-476), has been previously shown to interact with the MAD1^CTD^ (*41, 43–47*). AlphaFold (*48*) predicts this N-terminal α-helix to be separated by an unstructured intervening region from a second α-helix between residues 498-507 (**Fig. 3B**). Residues in this second α-helix are only conserved in metazoans, to which RZZ is limited. We generated a series of N-terminal deletion constructs derived from BUB1^437-521^ and tested them in solid-phase binding assays. For unclear reasons, BUB1^467-521^ interacted with RZZ *in vitro* even more robustly than BUB1^437-521^ (**Fig. 2G-H**). Truncation of ten additional amino acids (BUB1^477-521^) strongly decreased the binding affinity to RZZ, though some residual binding was still observed relative to GST negative control. Any further deletions, including from the C-terminus, nearly eliminated RZZ binding (**Fig. S3E-F**). The robust binding of RZZ to BUB1^467-521^ predicts that BUB1 phosphorylation is dispensable for this interaction. Indeed, phosphorylated and unphosphorylated BUB1^437-521^ showed similar binding affinities for RZZ (**Fig. S5E-F**).

**Fig. 3.**
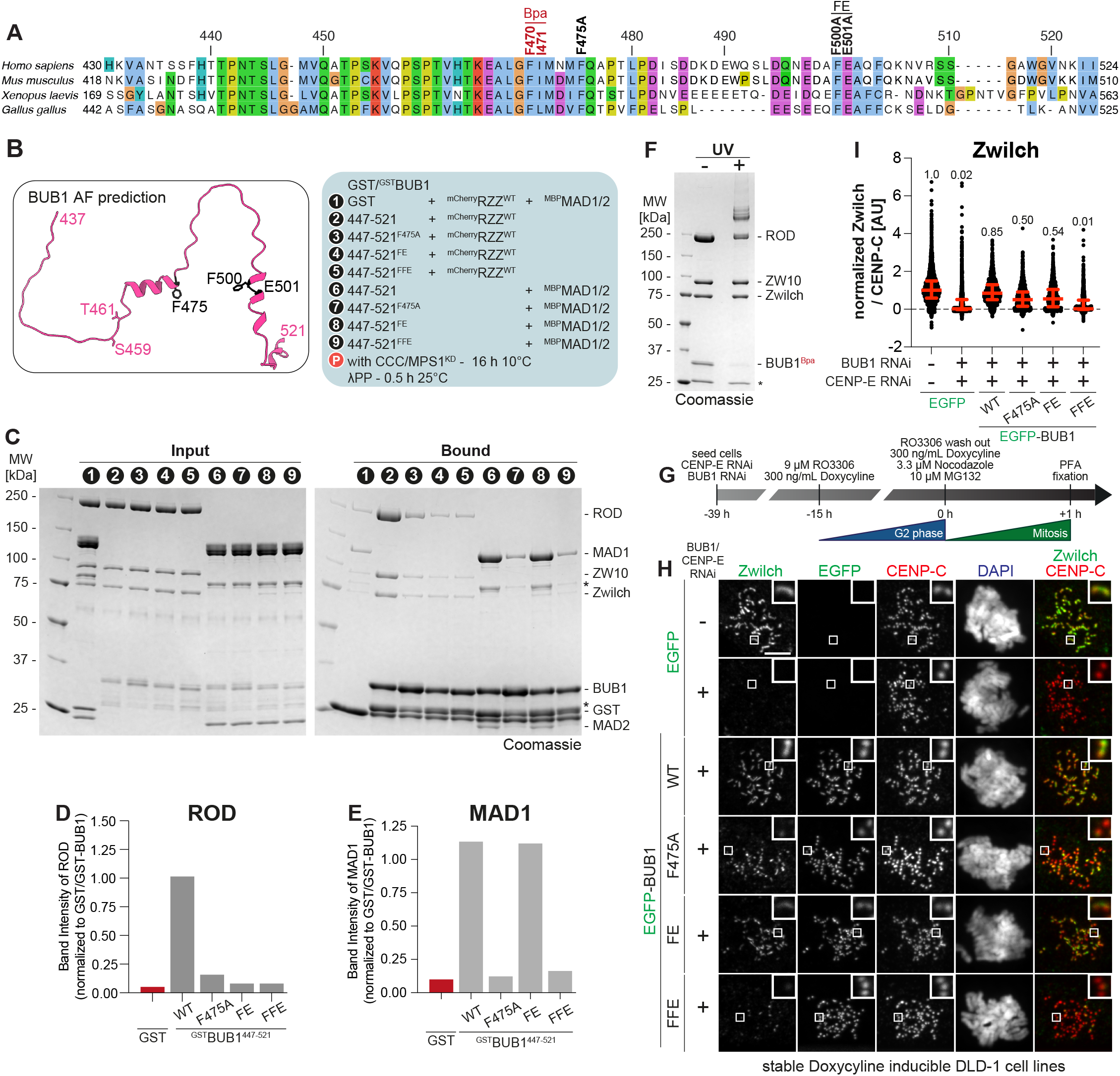
Validation of the RZZ-binding site of BUB1. (**A**) Multiple sequence alignment showing the central domain of BUB1. Mutations discussed in main text are indicated above the sequence alignment. (**B**) AlphaFold2 prediction of BUB1^437-521^. Amino acid substitutions are highlighted in black. (**C**) SDS-PAGE analysis of a pulldown assay with either GST or GST-tagged BUB1 baits, and mCherry-RZZ or MBP-MAD1/2 as prey. For overnight phosphorylation, bait and prey were incubated with CCC and MPS1^KD^ for 16 h at 10°C. Before SDS-PAGE analysis the eluate was dephosphorylated with λ-phosphatase for 30 mins at 25°C. (**D-E**) Quantification of the ROD and MAD1 band intensity normalized to the bait signal from the SDS-PAGE depicted in **c** from two individual repeats. (**F**) SDS-PAGE analysis of successful crosslinking of a BUB1^BPA^ mutant with mCherry-RZZ upon treatment with UV light. The experiment was performed once. (**G**) Scheme for the cell synchronization and RNAi treatment for the experiment shown in (H). (**H**) Representative images showing the localization of Zwilch in stable DLD-1 cell lines expressing different BUB1 constructs treated as indicated in (G). CENP-C was used to visualize kinetochores and DAPI to stain DNA. Scale bar: 5 μm. (**I**) Quantification of Zwilch levels at kinetochores of the experiment shown in (H). Red bars represent median (value shown above scatter dot plot) and interquartile range of normalized kinetochore intensity values. n refers to individually measured kinetochores (left to right: 2423, 2033, 1152, 1016, 946, 980) from three independent experiments.

We introduced mutations in GST-BUB1^467-521^ focussing on conserved and exposed residues (**Fig. 3A-B**). Mutating Phe475 to alanine strongly reduced RZZ (**Fig. S4A-B**). Mutating Phe500 and Glu501 (467−521^FE^) reduced RZZ binding to background levels, as also did combining these mutations with Phe475 (467−521^FFE^) (**Fig. S4A-B**). Also in analytical size-exclusion chromatography (SEC) experiments, where elution is determined by size and shape, all BUB1 constructs encompassing residues 467-521, including GST-BUB1^467-521^ and GST-BUB1^209-521^:BUB3 bound RZZ and co-eluted at a reduced elution volume. Conversely, mutants carrying F475A, F500A-E501A, or their combination prevented binding in solution (**Fig. S4C-H**). Despite being paralogous to BUB1:BUB3, BUBR1:BUB3 did not bind RZZ (**Fig. S4I**). Thus, the BUB1 binding site for RZZ is bipartite, with two binding motifs that are both necessary, but individually insufficient, for a robust binding interaction *in vitro*. Confirming the importance of the first helix for MAD1 binding, both the F475A and the FFE mutants of BUB1^447−521^, which share a dysfunctional first helix, showed little to no binding to MAD1:MAD2. The BUB1 FE mutant, however, had no impact on MAD1:MAD2 interaction, contrary to RZZ binding (**Fig. 3C-E**). In line with the crucial role of F475 in both MAD1 and RZZ binding, the previously described BUB1^RRK^ mutant, where F475 is mutated together with two neighbouring residues (M474R, F475R, Q476K) (*43*), reduced both MAD1:MAD2 and RZZ binding to levels similar to those in the negative control (**Fig. S5A-D**).

Thus, RZZ interacts directly with a segment of BUB1 that is more extended of, but partly overlapping with, the one interacting with the MAD1^CTD^. We used amber codon suppression technology (*49, 50*) to map the interaction of BUB1 with RZZ. Replacement of BUB1 residues F470 and I471 with the UV-photoactivatable crosslinker p-Benzoyl-L-phenylalanine (BPA) (**Fig. 3A**), followed by UV cross-linking, unequivocally indicated that BUB1^467-521^ interacts with ROD (**Fig. 3F**).

### Validation of the RZZ-binding site of BUB1

To validate the biological significance of these findings, we co-depleted endogenous BUB1 and CENP-E (**Fig. S6A-B**) in stable DLD-1 cell lines engineered to express various EGFP-BUB1 fusions, including BUB1^Δ467-521^, BUB1^FFE^, BUB1^F475A^, BUB1^FE^, and a construct containing only the BUB3 binding domain (GLEBS motif; BUB1^209-270^) (**Fig. 3G** and **Fig. S6C**). In absence of transgene expression, RZZ and MAD1 were lost from kinetochores, as expected, whereas the levels of BUBR1 were only partially reduced (**Fig. 3H-I** and **Fig. S6H-I**). Expression of BUB1^FL^ largely restored RZZ and MAD1 levels at prometaphase kinetochores (**Fig. 3H-I** and **Fig. S6D-F**). Instead, expression of BUB1^Δ467-521^, BUB1^FFE^, or BUB1^209-270^ in cells depleted of BUB1 and CENP-E completely abrogated kinetochore recruitment of RZZ and MAD1 (**Fig. 3H-I** and **Fig. S6D-J**), whereas BUB1^F475A^ and BUB1^FE^ had intermediate effects, in line with the biochemical experiments. Because the kinetochore levels of BUBR1 were fully restored in presence of BUB1^FFE^ (and of BUB1^F475A^, or BUB1^FE^) (**Fig. S6I**), our observations confirm that BUBR1 cannot replace BUB1 in RZZ or MAD1 recruitment when CENP-E is depleted.

### Two BUB1-dependent axes of corona assembly

Our studies identify two BUB1-dependent branches for RZZ recruitment, namely 1) a direct interaction of BUB1^467-521^ with RZZ, and 2) an interaction of BUB1 with BUBR1 that in turn recruits CENP-E for RZZ binding. If correct, this model predicts that a BUB1 mutant impaired in its ability to recruit BUBR1-CENP-E and deleted of the 467-521 region that binds RZZ ought to be completely unable to recruit the RZZ. Indeed, a BUB1 mutant lacking residues 271-409 and 467-521 (BUB1^Δ271-409/467-521^) caused complete elimination of RZZ from kinetochores (**Fig. 4A-F**), while deletion of only the BUB1 helix (BUB1^Δ271-409^) prevented BUBR1 recruitment but still supported robust RZZ localization (**Fig. 4A-F**). Kinetochore CENP-E was reduced in cells expressing BUB1^Δ271-409^, and further reduced in cells expressing BUB1^Δ271-409/467-521^, in agreement with the idea that BUBR1 and RZZS contribute to CENP-E recruitment.

**Fig. 4.**
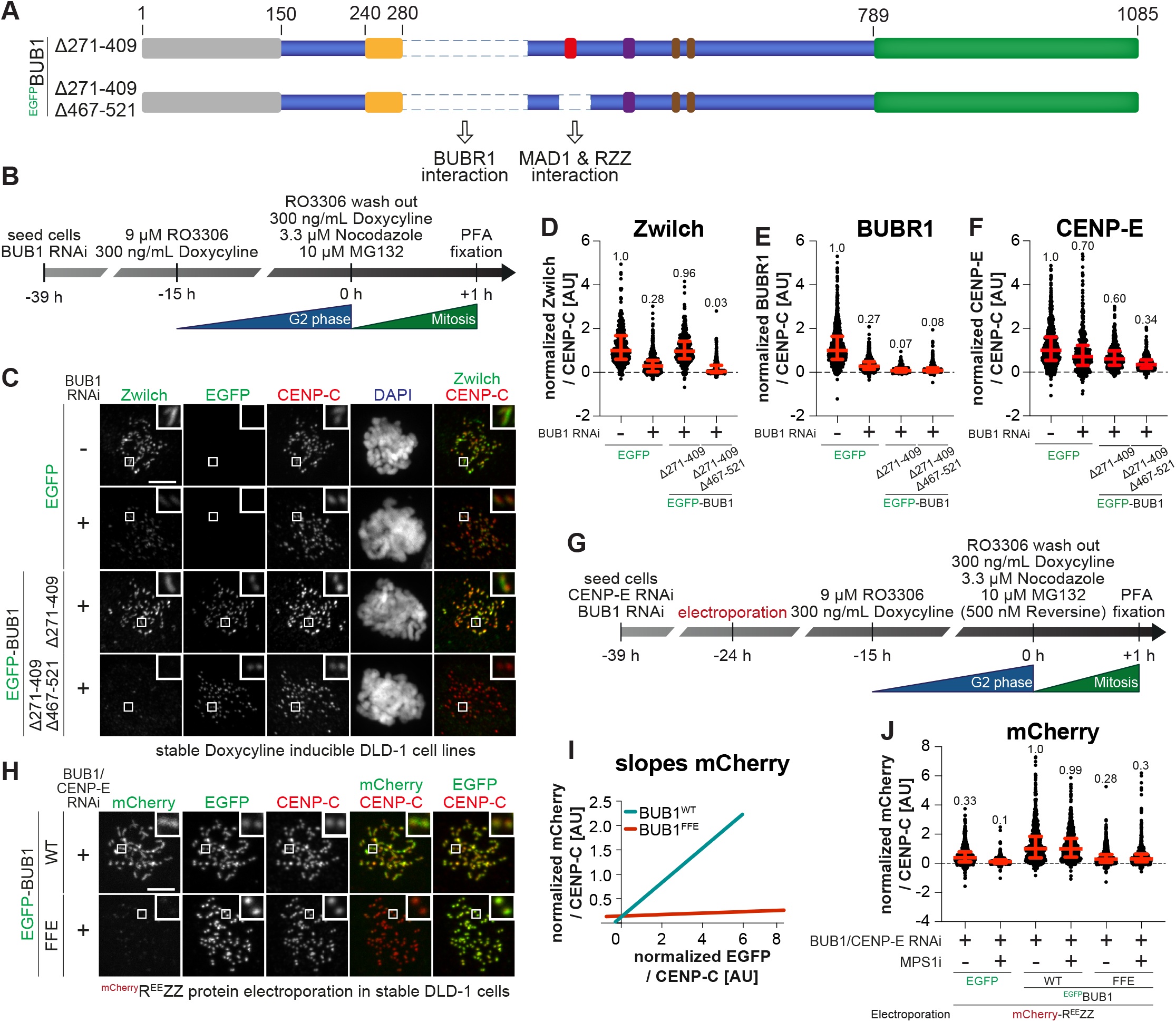
BUB1 and BUBR1 paralogues recruit RZZ to the kinetochore. (**A**) Scheme of the stable DLD-1 cell lines used in the experiment depicted in (C). Cells expressed N-terminally EGFP-tagged BUB1 constructs upon addition of doxycycline. (**B**) Scheme for cell synchronization and RNAi treatment for the experiment shown in (C). (**C**) Representative images showing the localization of Zwilch in stable DLD-1 cell lines expressing different BUB1 constructs treated as indicated in (B). CENP-C was used to visualize kinetochores and DAPI to stain DNA. Scale bar: 5 μm. (**D**) Quantification of Zwilch levels at kinetochores for experiment shown in (C). Red bars represent median (value shown above scatter dot plot) and interquartile range of normalized kinetochore intensity values. n refers to individually measured kinetochores (left to right: 304, 614, 362, 391). The experiment was performed three times. (**E**) Quantification of BUBR1 levels at kinetochores of the experiment shown in (C). Red bars represent median (value shown above scatter dot plot) and interquartile range of normalized kinetochore intensity values. n refers to individually measured kinetochores (left to right: 809, 825, 246, 464). The experiment was performed three times. (**F**) Quantification of CENP-E levels at kinetochores of the experiment shown in (C). Red bars represent median (value shown above scatter dot plot) and interquartile range of normalized kinetochore intensity values. n refers to individually measured kinetochores (left to right: 3672, 438, 504, 393). CENP-E staining was performed once. (**G**) Scheme of cell synchronization and RNAi treatment for the experiment shown in (H). (**H**) Representative images showing stable DLD-1 cell lines electroporated with mCherry-R^EE^ZZ according to the experimental scheme shown in (G). CENP-C was used to visualize kinetochores and DAPI to stain DNA. Scale bar: 5 μm. (**I**) Least-square linear fitting of data points for each kinetochore of the mCherry intensity on the y-axis and the EGFP intensity on the x-axis of the experiment shown in (H). (**J**) Quantification of mCherry levels at kinetochores of the experiment shown in (H). Red bars represent median (value shown above scatter dot plot) and interquartile range of normalized kinetochore intensity values. n refers to individually measured kinetochores (left to right: 569, 620, 632, 657, 792, 658). The experiment was performed twice.

To verify that the elusive kinetochore receptor of RZZ is indeed the binding site on BUB1 we have identified, we applied our stringent experiment with electroporated mCherry-tagged R^EE^ZZ in cells depleted of endogenous CENP-E and BUB1 and expressing either full-length BUB1^WT^ or BUB1^FFE^ transgenes (**Fig. 4G**). mCherry-R^EE^ZZ localized robustly to kinetochores in cells expressing BUB1^FL^, but only minimal residual localization was observed in cells expressing BUB1^FFE^ (**Fig. 4H-J**). Thus, when CENP-E depletion is coupled with mutations that prevent the interaction of BUB1 with RZZ, even constitutive polymerization in presence of endogenous MPS1 activity is insufficient for kinetochore recruitment of RZZS. Collectively, these findings provide a compelling recruitment model for the RZZ, as further discussed in Conclusions.

### Cooperative binding of MAD1 and the RZZ to BUB1

An unexpected conundrum of our study concerns the MAD1 recruitment mechanism, as we have shown that the MAD1^CTD^ and the RZZ compete for BUB1 binding. In principle, this problem may be solved by mobilizing a higher proportion of BUB1 than the sum of MAD1 and RZZ, thus buffering the effects of competition. This is a concrete possibility considering the relative cellular concentrations of these species and the fact that KNL1 exposes numerous MELT repeats to increase the multiplicity of BUB1 (*30, 51–57*). Nevertheless, a model purely based on competition seems unsatisfactory, as it would defy previous observations that RZZ facilitates MAD1 recruitment, rather than opposing it (*6, 26, 37, 39, 41, 42, 45, 58–63*). To shed further light on this question, we depleted BUB1 and CENP-E to eliminate MAD1 and RZZS. Re-expression of a BUB1^WT^ transgene largely restored kinetochore MAD1. Albeit at reduced levels, re-expression of BUB1^S459-T461A^, a mutant form of BUB1 that binds RZZ but not MAD1, also restored high levels of kinetochore MAD1 (**Fig. S7A-C**), confirming a role of RZZS in MAD1 recruitment downstream of BUB1.

To assess if RZZ binds MAD1:MAD2 directly, we polymerized RZZS into rings, as described previously (*9*) (**Fig. S7D**) and performed a pelleting assay with BUB1^467-521^ and full-length MAD1:MAD2. MAD1:MAD2 and BUB1^467-521^ pelleted with RZZS, while in absence of RZZS both proteins were almost entirely in the supernatant (**Fig. 5A**). As expected, MAD1^CTD^, which competes with RZZ for BUB1 binding, was instead found in the supernatant (**Fig. 5A**). In another set of experiments, we titrated increasing amounts of MAD1:MAD2 (0.5-8 µM) and RZZ (0.5-8 µM) in presence of BUB1 as bait (at 8 µM). We observed a stepwise increase in bound proteins until both components reached apparently equimolar concentrations (**Fig. 5B** and **Fig. S7E-F**). This experiment indicates unequivocally that despite the apparent competition of the MAD1^CTD^ and RZZ for BUB1, a MAD1 binding site on RZZ allows MAD1 to scale with RZZ on the BUB1 bait.

**Fig. 5.**
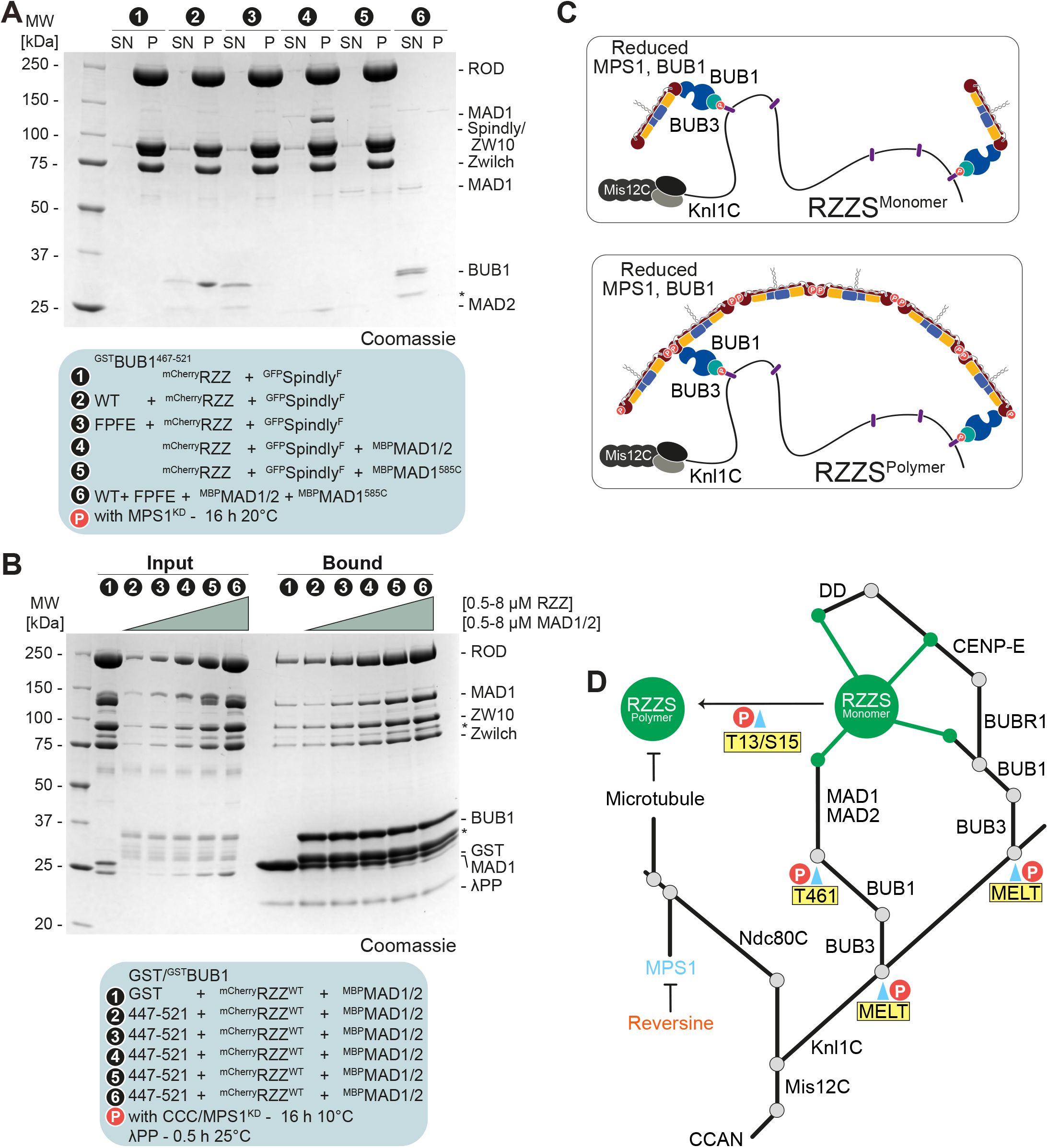
Cooperative binding of MAD1 and the RZZ to BUB1. (**A**) SDS-PAGE analysis of a pelleting assay. mCherry-RZZ and GFP-Spindly^F^ were polymerized overnight in presence of MPS1^KD^ at 20°C. Subsequently, polymerized RZZS was incubated with BUB1 or MAD1 constructs for an additional 2 h at 20°C and finally applied to a glycerol cushion and spun in an ultracentrifuge. (**B**) SDS-PAGE analysis of a pulldown assay with either GST or GST-tagged BUB1 as bait, mCherry-RZZ (concentrations indicated) and MBP-MAD1:MAD2 (concentrations indicated) as prey. Phosphorylation bait and prey were incubated with CCC (CDK1:Cyclin B:CKS1) and MPS1^KD^ for 16 h at 10°C. Prior to SDS-PAGE analysis the eluate was dephosphorylated with λ-phosphatase for 30 mins at 25°C. (**C**) Scheme depicting RZZ recruitment to the kinetochore through interaction with BUB1. (**D**) Revised hierarchal organization of outer kinetochore and kinetochore corona based on our new results.

### Conclusions

We have dissected the molecular basis of kinetochore recruitment of the kinetochore corona, an essential part of the kinetochore scaffold in early mitosis. The recruitment pathway has its basis in the BUB1:BUBR1 complex of paralogous proteins, whose functional specialization from an original singleton represents a stunning example of molecular evolution (*31, 33, 64*). The pathway consists of two branches. The first branch uses BUB1 as a direct receptor through a binding site on ROD. The second branch uses CENP-E, through BUBR1, which docks on BUB1 (*31, 36*). CENP-E interacts directly with an unknown binding site on the RZZ (*13*). While the two branches seem largely redundant for RZZS localization, we suspect them to be inseparable for proper corona function, as both contribute to robust localization of DD (*65*).

BUB1 lies at the fulcrum of the recruitment pathway. BUB1 is recruited largely through MPS1-dependent phosphorylation of the MELT repeats on KNL1, so that MPS1 inhibition reduces BUB1 kinetochore binding. R^EE^ZZ bypasses the requirement of MPS1 kinase for kinetochore localization when CENP-E has been depleted (*13*). Likely, this is not due to an increase in binding affinity of R^EE^ZZ for BUB1, as RZZ^WT^, R^AA^ZZ (the non-phosphorylatable mutant of RZZ), or R^EE^ZZ bound BUB1 with similar affinity in solid-phase or in solution (**Fig. S4H** and **Fig. S7G-H**). As R^EE^ZZ polymerizes when MPS1 is inhibited, its retention on CENP-E-depleted kinetochores indicates that it sticks more strongly than RZZ monomers to the residual BUB1. This is because monomers may be expected to require at least one corresponding BUB1 receptor for kinetochore recruitment, whereas a multivalent polymer may continue to bind even if only a handful of residual BUB1 remained after MPS1 inhibition (**Fig. 5C**). Thus, polymerization of the corona provides an extremely robust, cooperative mechanism to retain attachment to kinetochores, making it resilient to temporary changes in local phosphorylation patterns and to progressive depletion of binding sites as microtubule attachment proceeds. Residual BUB1 levels due to incomplete depletions, and/or ectopic binding of BUBR1/BUB3 to MELT repeats after depletion of BUB1 (*31, 33, 66*), both causing substantial residual RZZ docking, likely explain why previous analyses of the role of BUB1 in RZZ assembly in human cells led to conflicting conclusions (*26, 36–41*). Our observations also argue that BUB1, when bound to RZZ, is unlikely to satisfy a role as MAD1:MAD2 receptor. We surmise therefore that MAD1:MAD2 complexes recruited to RZZ interact with RZZ-free BUB1 molecules, which likely are numerous given the abundance of MELT repeats on KNL1 (**Fig. 5D**).

The physical interaction network of the corona integrates SAC and biorientation. By recruiting DD and CENP-E, the corona promotes transport of chromosomes to the spindle equator. By recruiting MAD1:MAD2, the corona promotes SAC signaling. The same mechanisms are also relevant for SAC silencing upon end-on conversion of microtubule attachment. This conversion detaches the chromosome cargo from the corona, activates DD, and allows corona disassembly, removing MAD1:MAD2 from the kinetochore and causing its suppression, which turns off the SAC. Suppression of MPS1 activity, which in turn responds to Aurora B kinase, a master regulator of biorientation, is key to SAC silencing. Progressive suppression of MPS1 causes progressive dephosphorylation of the MELT repeats and of MAD1:MAD2 binding to BUB1, and directly inactivates MAD1:MAD2 (*67, 68*). A crucial unresolved problem for future analyses is therefore how end-on conversion of microtubule attachment changes the inner fabric of interactions within the corona to allow DD activation and the transition to SAC silencing.

## Acknowledgements

We thank Sabine Wohlgemuth for protein purifications.

## Author contributions

Conceptualization: A.M., V.C.; Investigation: V.C.; Funding acquisition: A.M.; Project Administration: A.M.; Supervision: A.M. Validation: A.M., V.C.; Visualization: V.C., A.M; Writing – original draft: A.M., V.C.

## Data and materials availability

All data are available in the main text or the supplementary materials.

## Supplementary Materials

Materials and Methods

Figures S1 to S7

Tables S1 to S3

References for Supplementary

