## Supplementary Material for "The BUB1 and BUBR1 paralogs scaffold the kinetochore corona"

**The PDF file includes:**

Materials and Methods
Supplementary text
Figures S1 to S7
Tables S1 to S3
References for Supplementary

### Materials and Methods

Mutagenesis and cloning

The KNL1 constructs (isoform 2; UniProt H0YN41) were subcloned into pCDNA5/FRT/TO-EGFP-IRES as previously described (*1*). The human BUB1 sequence (UniProt O43683) was subcloned into a pCDNA5/FRT/TO-IRES plasmid with an N-terminal EGFP-tag to generate stable cells. The GST-tagged BUB1 truncations, used *in vitro*, were cloned into a pGEX vector with an N-terminal PreScission-cleavable GST-tag. For the $^{GST}$BUB1$^{209-521_MBP}$/BUB3 construct, the codon-optimized sequence was cloned into a pFLMultiBac vector (*2*). Plasmids for full-length BUB1/BUB3, BUBR1/BUB3, RZZ, Spindly and MAD1/2 were used as previously described (*3-6*). Mutations and deletions were introduced via site-directed mutagenesis and Gibson assembly (*7*) and verified by Sanger sequencing (Microsynth Seqlab).

Expression and purification of RZZ, Spindly and CCC

The RZZ and Spindly constructs were expressed and purified using the biGBac system (*8*), as previously described (*4, 5, 9*). Expression and purification of human CDK1:Cyclin-B:CKS1 (CCC) was carried out as previously described (*10*).

Expression and purification of $^{GST}$BUB1, $^{MBP}$MAD1$^{585C}$

$^{GST}$BUB1 constructs were expressed in *Escherichia coli* BL21 CodonPlus cells, which were grown in 1 L TB at 37°C to an $OD_{600}$ of 0.6. The expression was induced with 0.2 mM IPTG and the culture was transferred to an incubator pre-cooled to 18°C and grown overnight before harvesting. The pellet was then snap-frozen and stored at −80°C until purification. Expression of $^{MBP}$MAD1$^{585C}$ was carried out in insect cells. The baculovirus was generated in Sf9 cells and used to infect 500 mL of Tnao38 cells, which were grown for 72 h at 27°C. The pellet was then snap-frozen and stored at −80°C until purification. To start the purification the pellet was resuspended in lysis buffer (50 mM HEPES pH 8.0, 250 mM NaCl, 2 mM Tris(2-carboxyethyl)phosphine (TCEP)) supplemented with protease inhibitor, 1 mM PMSF, DNaseI and lysed by sonication. The lysate was clarified by centrifugation for 45 minutes at 88,000x g, sterile filtered and loaded onto a GSTrap column (Cytiva). Subsequently, the column was washed with at least 20 column volumes of lysis buffer. Elution was performed with lysis buffer supplemented with 25 mM GSH. The eluate was and concentrated and loaded onto a Superdex 200 16/60 pre-equilibrated in SEC buffer (50 mM HEPES pH 8.0, 250 mM NaCl, 2 mM TCEP). Peak fractions containing the protein of interest were analysed by SDS-PAGE, concentrated, snap-frozen, and stored at -80˚C until further usage.

Expression and purification of $^{GST}$BUB1$^{209-521_MBP}$/BUB3, BUB1/BUB3 and BUBR1/BUB3

$^{GST}$BUB1$^{209-521_MBP}$/BUB3, BUB1/BUB3 and BUBR1/BUB3 were expressed using the biGBac system as 6xHis fusions. The baculovirus was generated in Sf9 cells and used to infect 1 L of Sf9 cells, which were grown for 72 h at 27°C. Proteins were immediately purified after harvesting by centrifugation. The pellet was resuspended in lysis buffer (50 mM HEPES pH 8.0, 300 mM NaCl, 5 mM imidazole pH 8, 2 mM $MgCl_2$, 2 mM TCEP) supplemented with protease inhibitor, 1 mM

PMSF, DNaseI and lysed by sonication. The lysate was clarified by centrifugation for 45 minutes at 88,000x g, sterile filtered and loaded onto a HisTrap HP column (Cytiva). Subsequently, the column was washed with at least 20 column volumes lysis buffer. The elution was performed with lysis buffer supplemented with 300 mM imidazole. The eluate was diluted 1:10 in no salt buffer (50 mM HEPES pH 8.0, 2 mM $MgCl_2$, 2 mM TCEP), and applied to a 6 ml ResourceQ anion exchange column (Cytiva). The protein was eluted with a 50-500 mM NaCl gradient over 10 column volumes, analysed by SDS-PAGE and fractions containing the protein of interest were pooled and concentrated with a 100 kDa cutoff Amicon concentrator (Millipore). After three buffer exchanges with SEC buffer (50 mM HEPES pH 8.0, 150 mM NaCl, 2 mM $MgCl_2$, 1 mM TCEP) the protein was concentrated to the desired concentration, snap-frozen, and stored at -80˚ C until further usage.

Expression and purification of $^{MBP}$MAD1/2$^{FL}$

Expression of $^{MBP}$MAD1/2$^{FL}$ was carried out in insect cells. The Baculoviruses were generated in Sf9 cells and for protein expression Tnao38 were co-infected with $^{MBP}$MAD1 and MAD2$^{6His}$ cells for 72 hours at 27°C before harvesting. The pellet was washed with PBS, snap-frozen and stored at -80˚ C. For purification, the MAD1/2$^{FL}$ pellet was resuspended in lysis buffer (50 mM HEPES pH 8.0, 250 mM NaCl, 15 mM imidazole pH 8, and 1 mM TCEP) supplemented with protease inhibitor, 1 mM PMSF, DNaseI and lysed by sonication. The lysate was clarified by centrifugation for 45 minutes at 88,000x g, sterile filtered and loaded onto a HisTrap HP column (Cytiva). Elution was performed with lysis buffer supplemented with 300 mM imidazole. Subsequently, the eluate was diluted 1:10 in zero salt buffer (50 mM HEPES pH 8.0, 1 mM TCEP), and applied to a 6 ml ResourceQ anion exchange column (Cytiva). The protein was eluted with a 50-500 mM NaCl gradient over 10 column volumes, analysed by SDS-PAGE and fractions containing the protein of interest were pooled and concentrated with a 100 kDa cutoff Amicon concentrator (Millipore). After three buffer exchanges with SEC buffer (50 mM HEPES pH 8.0, 250 mM NaCl, 1 mM TCEP) the protein was concentrated, snap-frozen, and stored at -80˚C until further usage.

Expression and purification of $^{mCherry}$MPS1$^{KD}$

Expression of the MPS1 kinase domain (KD) was carried out in insect cells as a 6xHis fusion protein. The baculovirus was generated in Sf9 cells and used to infect 500 mL of Tnao38 cells, which were grown for 72 h at 27°C. The pellet was washed with PBS, snap-frozen and stored at -80˚ C. For purification, the pellet was resuspended in lysis buffer (50 mM HEPES pH 8.0, 300 mM NaCl, 5% glycerol, 20 mM imidazole pH 8, and 2 mM TCEP) supplemented with protease inhibitor, 1 mM PMSF, DNaseI and lysed by sonication. The lysate was clarified by centrifugation for 45 minutes at 88,000x g, sterile filtered and loaded onto a HisTrap HP column (Cytiva). Elution was performed with lysis buffer supplemented with 300 mM imidazole. The eluate was concentrated and loaded onto a Superdex 200 10/300 pre-equilibrated in SEC buffer (50 mM HEPES pH 8.0, 300 mM NaCl, 5% glycerol, and 1 mM TCEP). Peak fractions containing the protein of interest were analysed by SDS-PAGE, concentrated, snap-frozen, and stored at -80˚ C until further usage.

Pulldown assays

For pulldown experiments, GST and GST-BUB1 or farnesylated GST-Spindly baits were mixed with preys at the indicated concentrations in 40 µl binding buffer (50 mM HEPES pH 8.0, 100 mM NaCl, 1 mM TCEP) and spun at 4°C for 30 minutes at 20,000x g. Where indicated phosphorylation was carried out by incubating prey and bait for 16 h at 10°C with $^{mCherry}$MPS1$^{KD}$ (1:20 kinase to substrate ratio) and CCC (1:100 kinase to substrate ratio) in binding buffer supplemented with 2 mM ATP, 5 mM $MgCl_2$ followed by a centrifugation step at 4°C for 30 minutes at 20,000x g. Subsequently, 3 µl of input was taken and mixed with 3 µl of 5xSDS-buffer. The remaining 35 µl of the protein mixtures were added to 10 µl dried GSH beads (pre-equilibrated in binding buffer) in Pierce micro-spin columns (Thermo Fischer Scientific) and incubated for 2 h at 4°C 300-rpm shaking. After the incubation step unbound protein was removed by centrifugation (2 minuets, 800x g, 4°C) and washed three times with 200 µl of binding buffer with a centrifugating step in-between (2 minuets, 800x g, 4°C). Finally, the bound fraction was incubated in 20 µl binding buffer supplemented with 50 mM glutathione pH 8 for 10 minuets and eluted by centrifugation (2 minuets, 1000 g, 4°C). Where indicated the eluate was treated for 30 mins with λ-phosphatase (produced in house) at 25°C. Finally, 5 µl of 5x SDS-sample buffer were added to the eluate and analysed via SDS-PAGE and Coomassie Blue staining or in-gel fluorescence, respectively.

Analytical Size Exclusion Chromatography

Binding assays in solution were performed under isocratic conditions on a Superose 6 15/50 pre-equilibrated in SEC buffer (50 mM HEPES pH 8.0, 100 mM NaCl, 1 mM TCEP) at 4°C on an ÄKTA pure micro system. The protein absorbance was monitored at 280 nm and the proteins were eluted in 100 µl fractions and analysed by SDS-PAGE and, Coomassie Blue staining. To assess complex formation, proteins were mixed at the indicated concentrations in 60 µl SEC buffer and incubated for 16 h at 10°C. Phosphorylation was carried out as indicated in the figure.

RZZ-Spindly ring formation and pelleting assay

RZZS filaments were formed as described in (*4*). Briefly, 4 µM $^{mCherry}$RZZ and 8 µM farnesylated $^{GFP}$Spindly (*5*) were incubated for 16 h at 20°C in presence of 1 µM $^{mCherry}$MPS1$^{KD}$ in 20 µl assay buffer (50 mM HEPES pH 8, 100 mM NaCl, 2 mM $MgCl_2$, 2 mM ATP, 1 mM TCEP). Subsequently, the preys were added for another 2 h at 20°C. Next, the protein mixtures were carefully added on 100 µl glycerol cushion (50 mM HEPES pH 8, 100 mM NaCl, 2 mM $MgCl_2$, 1 mM TCEP, 40% glycerol) and centrifuged at 100,000x g for 45 minuets at 20°C. After spinning, the supernatant was carefully removed and added to 5x SDS-buffer, the glycerol cushion was discarded, and the pellet resuspended in 5x SDS-buffer. Finally, supernatant and pellet were analysed by SDS-PAGE followed by Coomassie Blue staining and measurement of the in-gel fluorescence.

Molecular modelling

BUB1 predictions were generated using AlphaFold Multimer (*11, 12*). Structures and predictions were analysed using ChimeraX-1.5 (*13*).

Cell culture and generation of stable cell lines

HeLa and DLD-1 were grown in Dulbecco's Modified Eagle's Medium (DMEM; PAN Biotech) supplemented with 10% tetracycline-free FBS (PAN Biotech), and L-Glutamine (PAN Biotech). Cells were grown at 37°C in presence of 5% $CO_2$. Parental Flp-In T-REx DLD-1 osTIR1 cells were a kind gift from D. C. Cleveland (University of California, San Diego, USA). Stable DLD-1 cell lines were generated using FRT/Flp recombination. KNL1 and BUB1 constructs were cloned into a pCDNA5/FRT/TO-EGFP-IRES plasmid and co-transfected with pOG44 (Invitrogen), encoding the Flp recombinase, into cells using X-tremeGENE (Roche) according to the manufacturer's instructions. Subsequently, cells were selected for 2 weeks in DMEM supplemented with hygromycin B (250 µg/ml; Thermo Fisher Scientific) and blasticidin (4 µg/ml; Thermo Fisher Scientific). Single-cell colonies were isolated, expanded and transgeme expression was induced by addition of 0.3 µg/ml doxycycline (Sigma-Aldrich) and checked by immunofluorescence microscopy and immunoblotting.

RNAi and drug treatment

Depletion of endogenous proteins was achieved through transfection of single small interfering RNA (siRNA) using RNAiMAX (Invitrogen) according to manufacturer's instructions. BUB1, CENP-E and KNL1 were depleted for 36 h through single transfections. Ndc80C was depleted for 48 h through two transfections. For a complete list of siRNA oligos used in this study see **Table S1**. Unless indicated otherwise, nocodazole (Sigma-Aldrich) was used at 3.3 µM, RO3306 (Calbiochem) at 9 µM, MG-132 (Calbiochem) at 10 µM, and Reversine (Cayman Chem.) at 500 nM.

Electroporation of recombinant protein into human cells

Recombinant $^{mCherry}$RZZ, constructs were electroporated as previously described (*14*) using the Neon Transfection System Kit (Thermo Fisher Scientific). HeLa cells were depleted of the indicted proteins as outlined above. Subsequently, cells were trypsinized, washed with PBS, and resuspended in electroporation buffer R (Thermo Fisher Scientific). Recombinant protein was added at a final concentration of 7 µM and electroporated by applying two consecutive 35 ms pulses with an amplitude of 1000 V (HeLa) or 1 pulse for 40 ms with an amplitude of 1400 V (DLD-1). Control cells were electroporated with mCherry or electroporation buffer, respectively. The electroporated sample was subsequently added to 15 ml of prewarmed PBS, centrifuged at 1500 rpm for 5 minutes, and incubated with Trypsin/ EDTA for 5 minutes. After two PBS washes, the cell pellet was resuspended in prewarmed DMEM and seeded in a 6-well plate with poly-L-lysine coated coverslips. Following an 8 h recovery, cells were treated with 9 µM RO3306 (Calbiochem) for 15 h and doxycycline (300 ng/mL) where indicated. Subsequently, cells were released into mitosis in presence of 3.3 µM nocodazole and 10 µM MG-132 for 1 h before fixation for immunofluorescence.

Immunofluorescence

IF experiments were carried out as previously described (*15*). Briefly, cells were grown on coverslips coated with poly-L-lysine (Sigma-Aldrich). Before fixation, cells were permeabilized with 0.5% Triton X-100 in PHEM (Pipes, HEPES, EGTA, $MgSO_4$) buffer supplemented with 100 nM microcystin for 5 minutes and fixed with 4% PFA in PHEM for 20 minutes. Following fixation cells were blocked with 5% boiled goat serum (BGS) in PHEM buffer for 1 h and subsequently incubated for 2 hours at room temperature with the respective primary antibodies in 2.5% BGS-PHEM supplemented with 0.1% Triton-X-100. For a complete list of primary antibodies used in this study see **Table S2**. Subsequently, cells were incubated for 1 hour at room temperature with the respective secondary antibodies (all 1:200 in 2.5% BGS-PHEM supplemented with 0.1% Triton-X-100). For a complete list of secondary antibodies used in this study see **Table S3**. All washing steps were performed with PHEM supplemented with 0.1% Triton-X-100 (PHEM-T) buffer. DNA was stained with 0.5 µg/ml DAPI (Serva) and Mowiol (Calbiochem) was used as mounting media.

Cell imaging

Cells were imaged at room temperature using a spinning disk confocal device on the 3i Marianas system equipped with an Axio Observer Z1 microscope (Zeiss), a CSU-X1 confocal scanner unit (Yokogawa Electric Corporation, Tokyo, Japan), 100 × /1.4NA Oil Objectives (Zeiss), and Orca Flash 4.0 sCMOS Camera (Hamamatsu). Confocal images were acquired as z-sections of 0.27 µm (using Slidebook Software 6 from Intelligent Imaging Innovations). Images were converted into maximal intensity projections, converted into 16-bit TIFF files, and exported. Automatic quantification of single kinetochore signals was performed using the software Fiji with background subtraction. Measurements were exported in Excel (Microsoft) and graphed with GraphPad Prism 10 (GraphPad Software). Figures were arranged using Adobe Illustrator 2025.

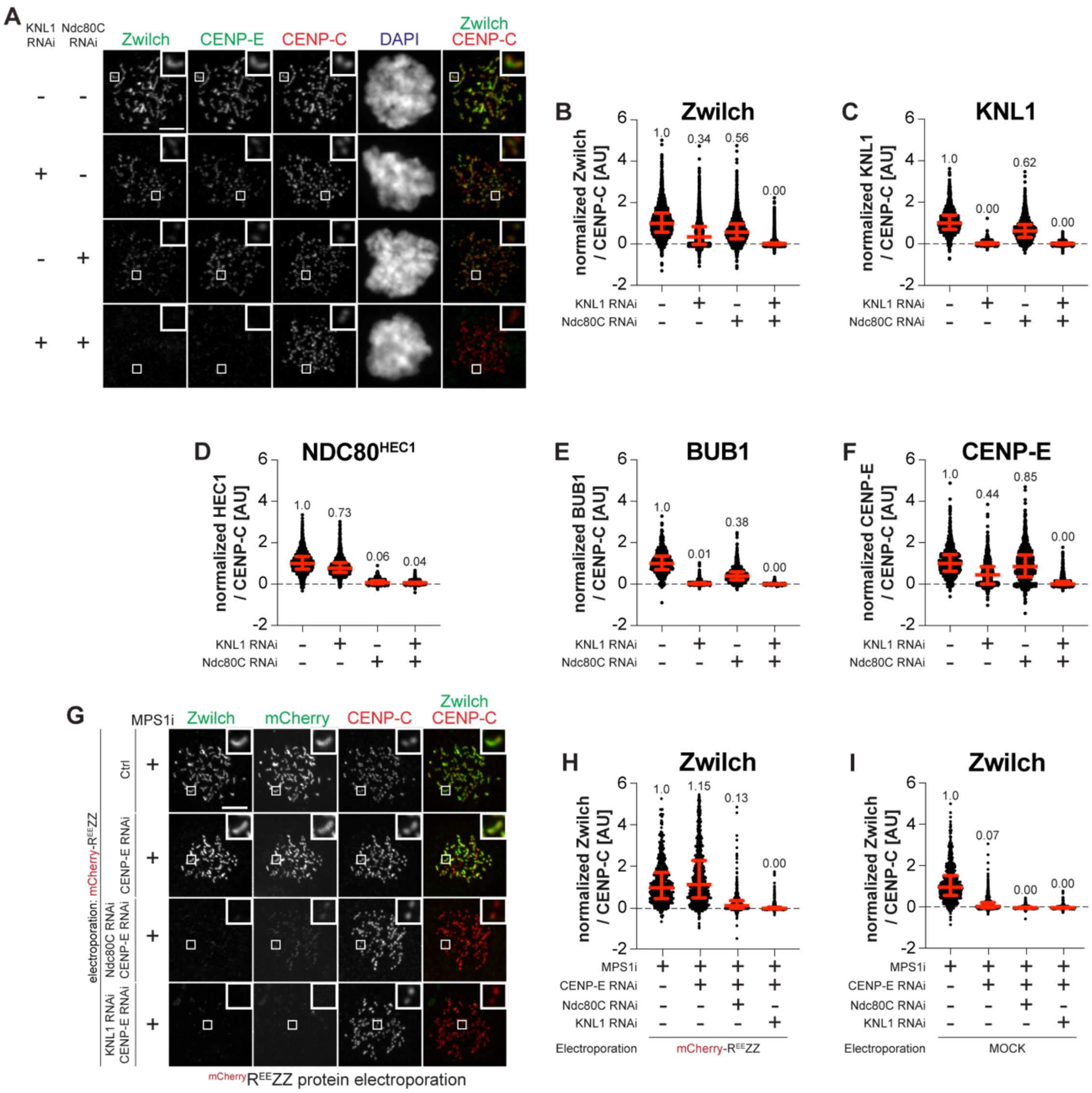


Supplement 1

**Fig. S1.** (**A**) Representative images of HeLa cells after RNAi treatment to deplete KNL1 and/or Ndc80C. NDC80C RNAi treatment was performed for 48 h. KNL1 RNAi was performed for 40 h. 8 h after the second transfection with siRNA to deplete Ndc80C, cells were synchronized in G2 phase with RO3306 for 15 h and then released into mitosis. Subsequently, cells were immediately treated with 3.3 µM nocodazole and 10 µM MG132 for an additional hour. CENP-C was used as a kinetochore marker and DAPI to visualize DNA. Scale bar: 5 µm (**B-F**) Quantification of Zwilch, KNL1, NDC80$^{HEC1}$, BUB1 and CENP-E levels at kinetochores of the experiment shown in (A). Red bars represent median (value shown above scatter dot plot) and interquartile range of normalized kinetochore intensity values. n refers to individually measured kinetochores (B left to right: 2369, 2557, 2670, 3381, C left to right: 1781, 1475, 1212, 1156, D left to right: 1781, 1475, 1212, 1156, E left to right: 638, 999, 815, 649, F left to right: 899, 1065,

1278, 1371) from three independent experiments. (**G**) Representative images showing the localization of Zwilch in cells electroporated with mCherry-$R^{EE}$ZZ treated with the respective RNAi as indicated in the figure according to the scheme shown in **Fig. 1D**. CENP-C was used to visualize kinetochores. Scale bar: 5 µm (**H-I**) Quantification of Zwilch levels at kinetochores of the experiment shown in (G). Red bars represent median (value shown above scatter dot plot) and interquartile range of normalized kinetochore intensity values. n refers to individually measured kinetochores (H left to right: 541, 1139, 469, 494, I left to right: 714, 1165, 702, 665) from two independent experiments.

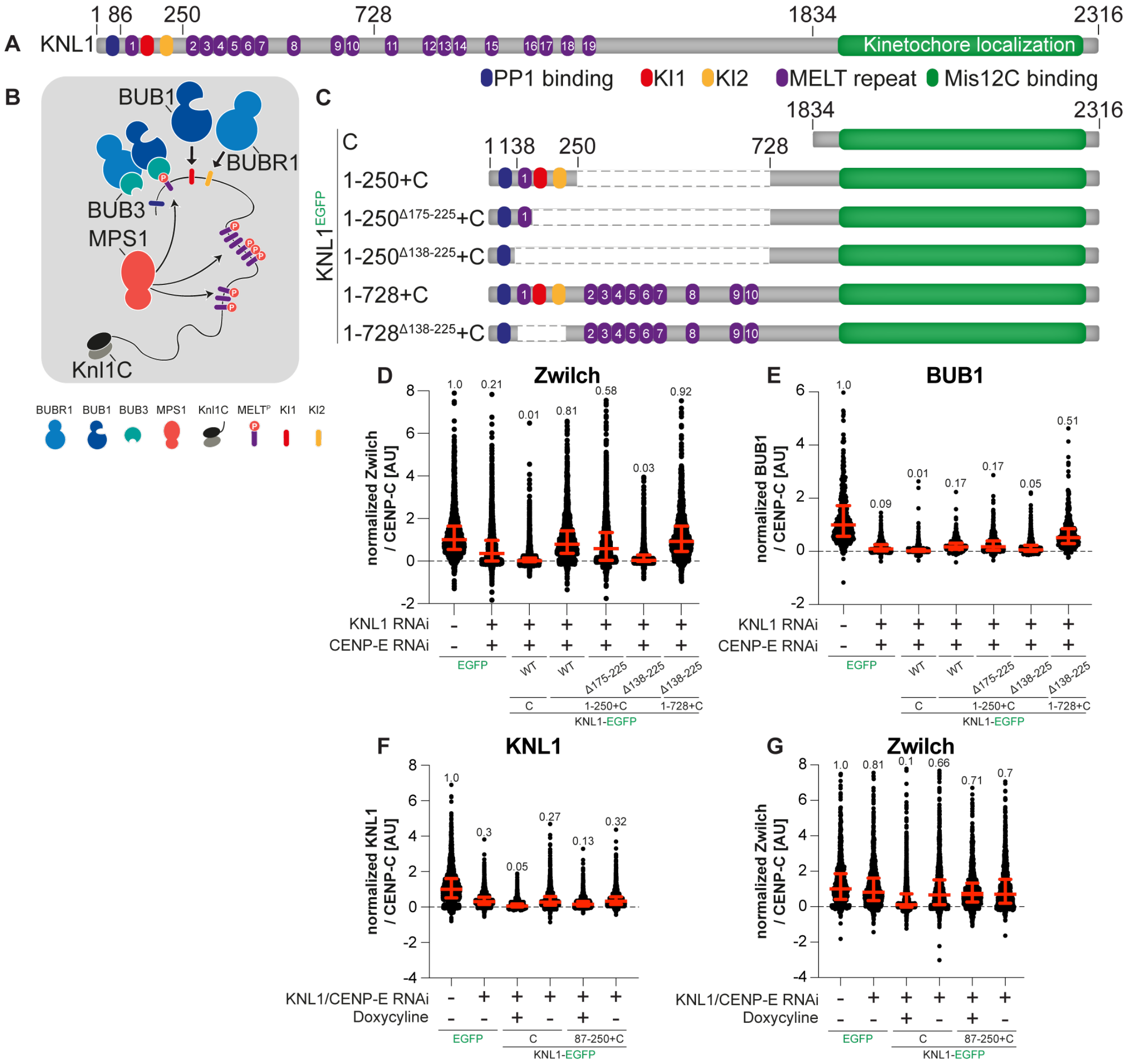


Supplement 2

**Fig. S2.** (**A**) Schematic representation of the organization of KNL1 with relevant functional domains. (**B**) Schematic depiction of the KNL mediated recruitment of BUB1:BUB3 and BUBR1:BUB3 through phosphorylated MELT repeats and the KI motifs. (**C**) Schematic of the stable DLD-1 cell lines used in the experiment depicted in **Fig. 1I**. Cells express C-terminally EGFP-tagged KNL1 constructs upon addition of doxycycline. (**D**) Quantification of Zwilch levels at kinetochores of the experiment shown in **Fig. 1I**. Red bars represent median (value shown above scatter dot plot) and interquartile range of normalized kinetochore intensity values. n refers to individually measured kinetochores (left to right: 1576, 2230, 2556, 1772, 1951, 1272) from three independent experiments. (**E**) Quantification of BUB1 levels at kinetochores of the experiment shown in **Fig. 1I**. Red bars represent median (value shown above scatter dot plot) and interquartile range of normalized kinetochore intensity values. n refers to individually measured kinetochores (left to right: 500, 823, 520, 514, 609, 537, 446) from three independent experiments. (**F**) Quantification of KNL1 levels at kinetochores of stable DLD-1 cell lines expressing different KNL1 constructs. Cells were depleted of KNL1 and CENP-E under suboptimal silencing conditions. 16 h before fixation protein expression was induced through addition of doxycycline and cells were synchronized in G2 phase with RO3306 for 15 h. Subsequently, cells were released into mitosis and immediately treated with 3.3 μM nocodazole, 10 μM MG132 and doxycycline for one hour before fixation. Red bars represent median (value shown above scatter dot plot) and interquartile range of normalized kinetochore intensity values. n refers to individually measured kinetochores (left to right: 1695, 2409, 2366, 2096, 1600, 2248) from three independent experiments. (**G**) Quantification of Zwilch levels at kinetochores of stable DLD-1 cell lines expressing different KNL1 constructs. Cells were depleted for KNL1 and CENP-E under suboptimal conditions. 16 h before fixation protein expression was induced through addition of doxycycline and cells were synchronized in G2 phase with RO3306 for 15 h. Subsequently, cells were released into mitosis and immediately treated with 3.3 μM nocodazole, 10 μM MG132 and doxycycline for one hour before fixation. Red bars represent median (value shown above scatter dot plot) and interquartile range of normalized kinetochore intensity values. n refers to individually measured kinetochores (left to right: 850, 979, 1575, 1485, 955, 1094) from three independent experiments.

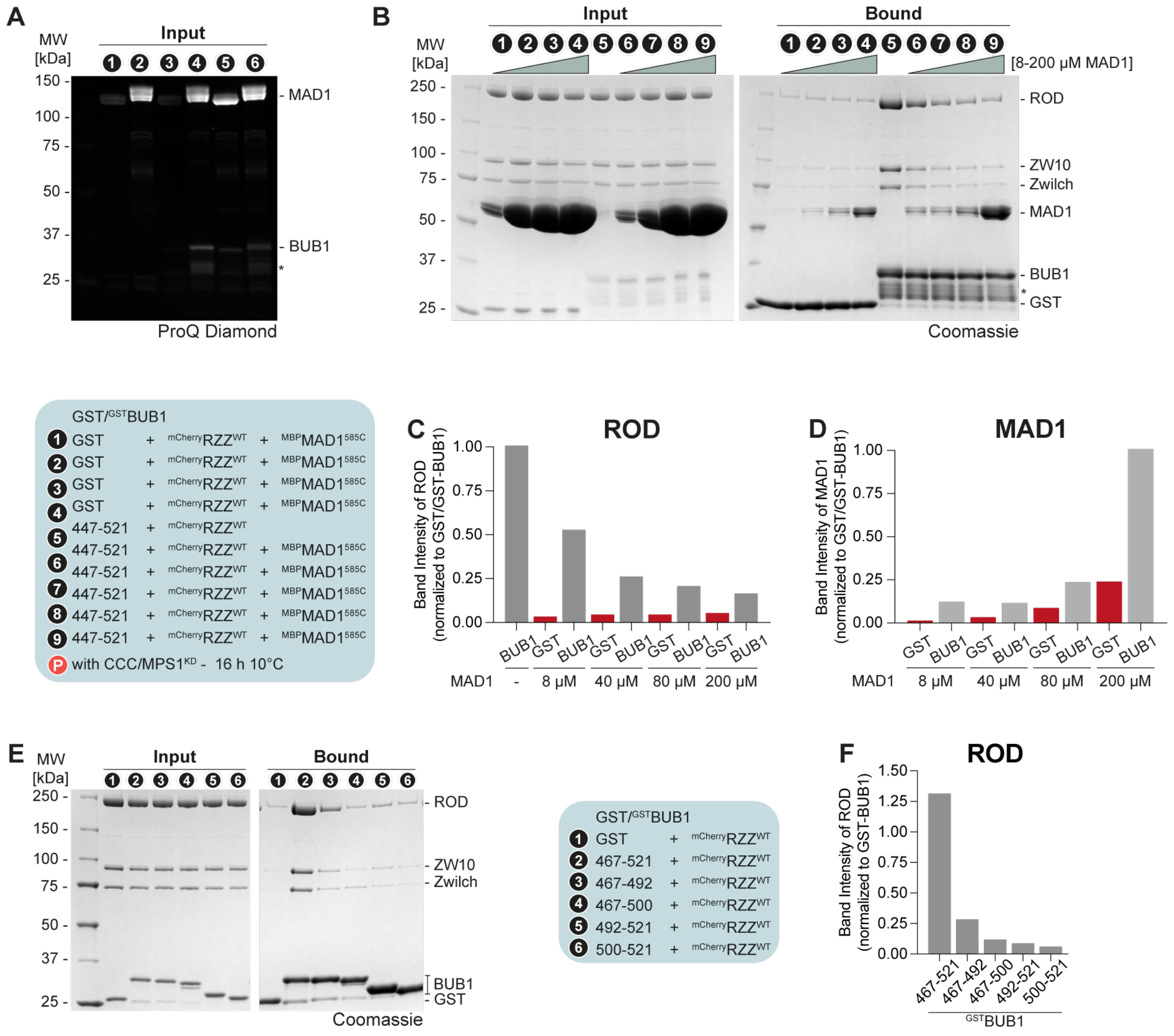


Supplement 3

**Fig. S3.** (**A**) ProQ Diamond staining was used to verify the successful phosphorylation of bait and prey from the experiment shown in **Fig. 2E**. (**B**) SDS-PAGE analysis of a pulldown assay with either GST or GST-tagged BUB1 as bait, mCherry-RZZ (concentrations indicated) and MBP-MAD1$^{585C}$ as prey (concentrations indicated). For overnight phosphorylation, bait and prey were incubated with CCC and MPS1$^{KD}$ for 16 h at 10°C. (**C-D**) Quantification of the ROD and MAD1 band intensity normalized to the bait signal from the SDS-PAGE depicted in (B). The experiment was repeated twice. (**E**) SDS-PAGE analysis of a pulldown assay with either GST or GST-tagged BUB1 constructs as bait, and mCherry-RZZ as prey. (**F**) Quantification of the ROD band intensity normalized to the bait signal from the SDS-PAGE depicted in (E). The experiment has been performed three times.

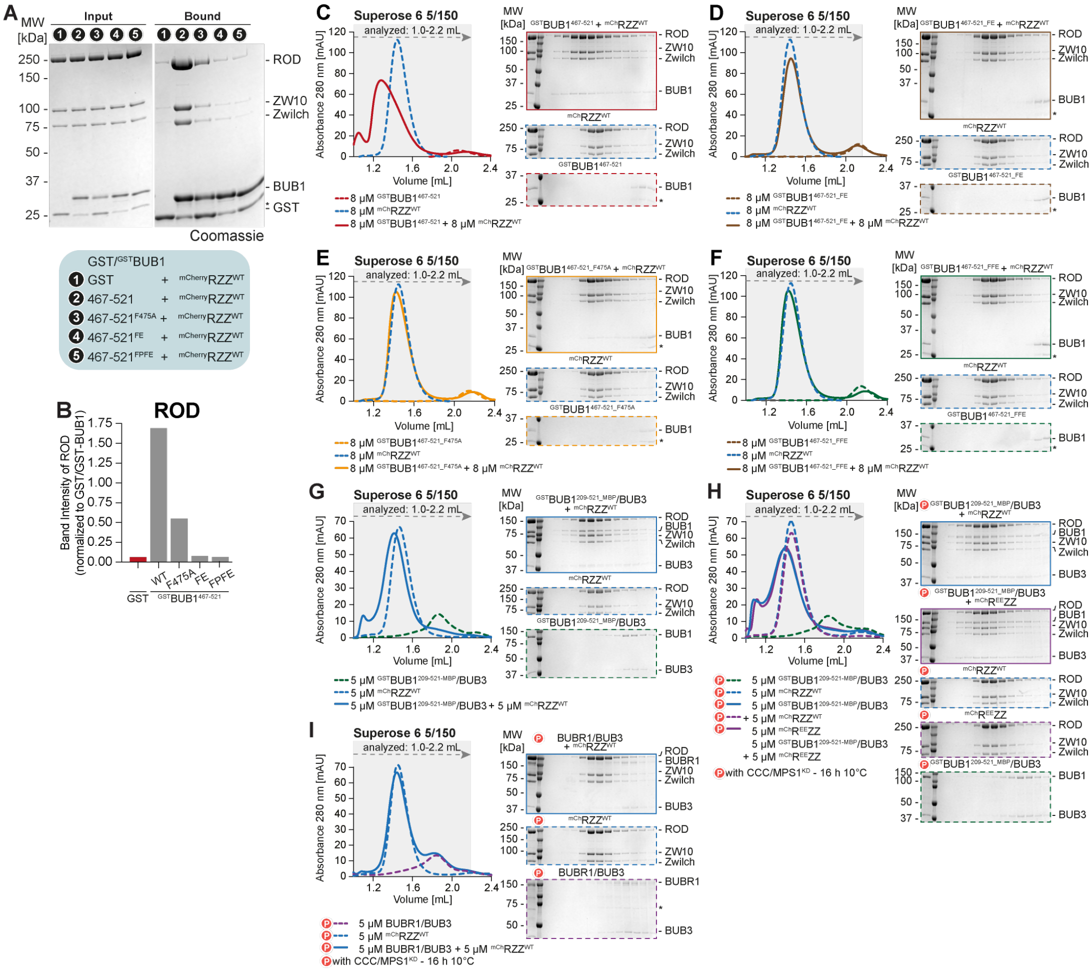


**Fig. S4.** (**A**) SDS-PAGE analysis of a pulldown assay with either GST or GST-tagged BUB1 constructs as bait and mCherry-RZZ as prey. (**B**) Quantification of the ROD band intensity normalized to the bait signal from the SDS-PAGE depicted in (A). (**C-I**) Analytical SEC binding assays with GST-BUB1 constructs, mCherry-RZZ constructs, and BUBR1/BUB3. The run with combined species is represented as a continuous line, and runs of individual species with a dashed line. The control gels with mCherry-RZZ alone are shared between panels (C-F). For overnight phosphorylation bait and prey were incubated with CCC and MPS1$^{KD}$ for 16 h at 10°C.

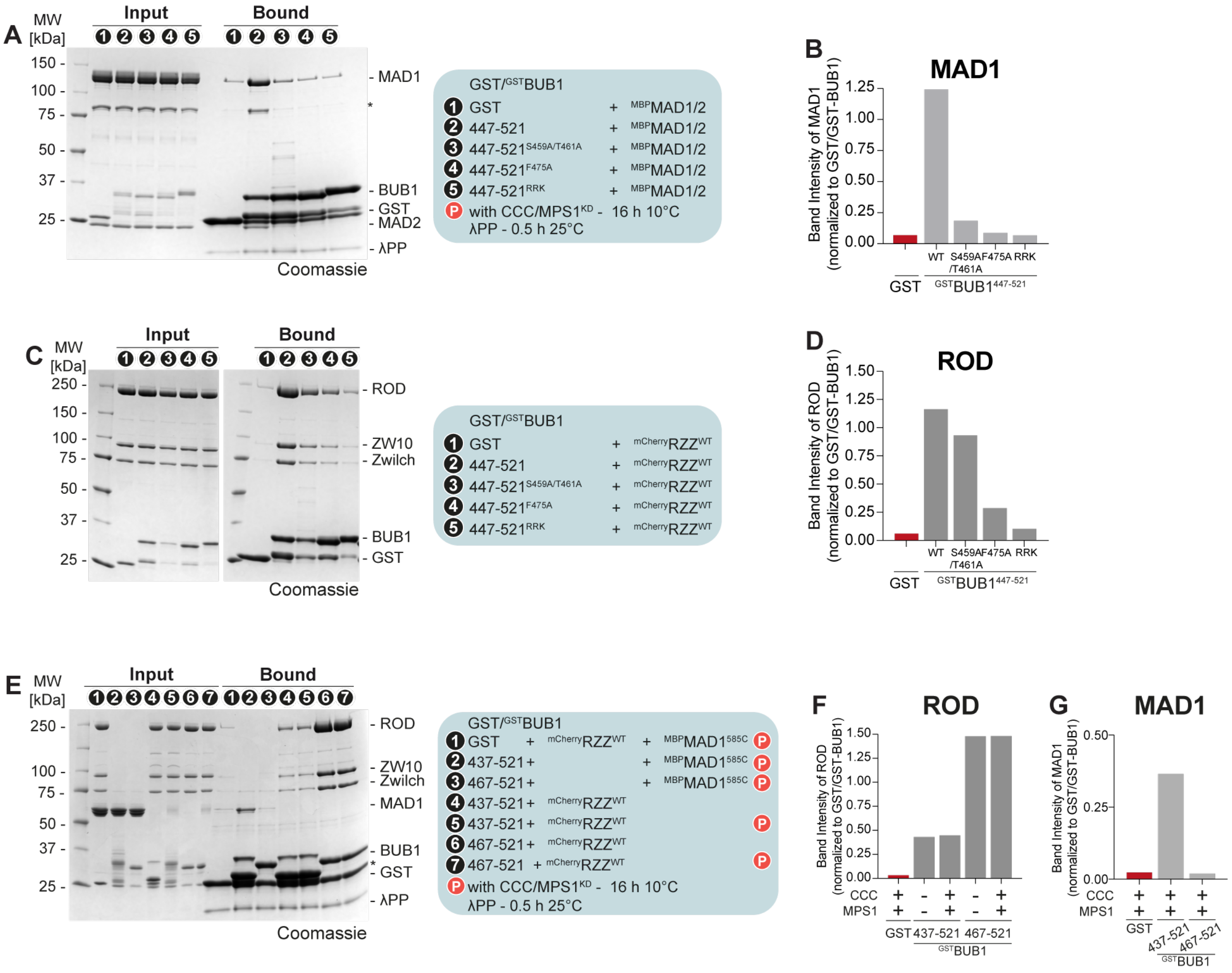


Supplement 5

**Fig. S5.** (**A**) SDS-PAGE analysis of a pulldown assay with either GST or GST-tagged BUB1 constructs as bait and MBP-MAD1:MAD2 as prey. For overnight phosphorylation bait and prey were incubated with CCC and MPS1$^{KD}$ for 16 h at 10˚C. Before SDS-PAGE analysis the eluate was dephosphorylated with λ-phosphatase for 30 mins at 25°C. (**B**) Quantification of the MAD1 band intensity normalized to the bait signal from the SDS-PAGE depicted in (A). The experiment was performed three times. (**C**) SDS-PAGE analysis of a pulldown assay with either GST or GST-tagged BUB1 constructs as bait and mCherry-RZZ as prey. (**D**) Quantification of the ROD band intensity normalized to the bait signal from the SDS-PAGE depicted in **c**. The experiment was performed three times. (**E**) SDS-PAGE analysis of a pulldown assay with either GST or GST-tagged BUB1 constructs as bait and mCherry-RZZ or MBP-MAD1/2 as prey. For overnight phosphorylation bait and prey were incubated with CCC and MPS1$^{KD}$ for 16 h at 10˚C. Before SDS-PAGE analysis the eluate was dephosphorylated with λ-phosphatase for 30 mins at 25°C. (**F-G**) Quantification of the ROD/MAD1 band intensity normalized to the bait signal from the SDS-PAGE depicted in (E). The experiment was performed once.

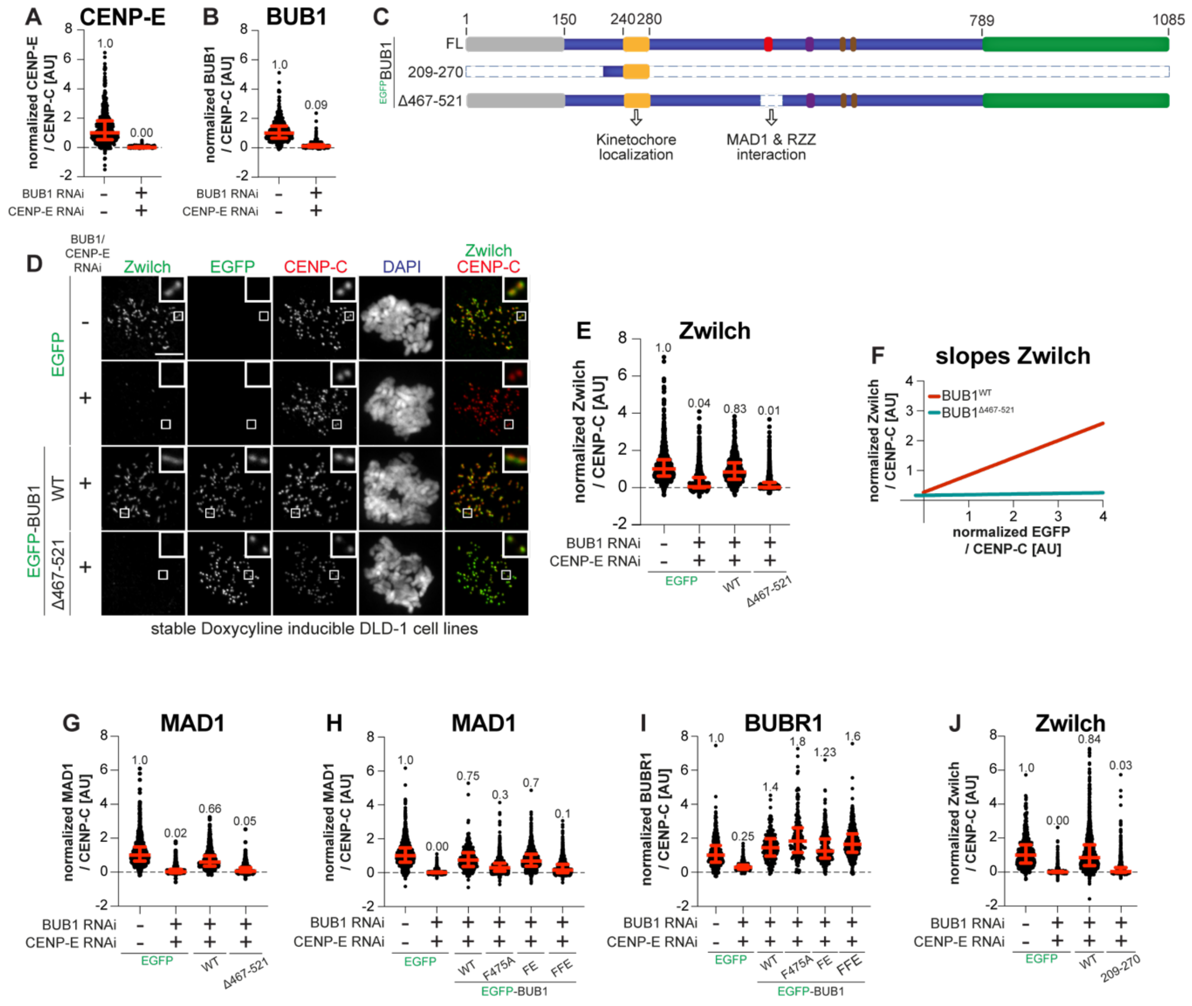


Supplement 6

**Fig. S6.** (**A-B**) Quantification of CENP-E and BUB1 levels at kinetochores of the experiment shown in **Fig. 3H**. Red bars represent median (value shown above scatter dot plot) and interquartile range of normalized kinetochore intensity values. n refers to individually measured kinetochores (A left to right: 633, 892, B left to right: 734, 949) (**C**) Schematic of the stable DLD-1 cell lines used in the experiment depicted in (D). Cells express N-terminally EGFP-tagged BUB1 constructs upon addition of doxycycline. (**D**) Representative images showing the localization of Zwilch in stable DLD-1 cell lines expressing different BUB1 constructs treated as indicated in **Fig. 3G**. CENP-C was used to visualize kinetochores and DAPI to stain DNA. Scale bar: 5 µm. (**E**) Quantification of Zwilch levels at kinetochores of the experiment shown in (D). Red bars represent median (value shown above scatter dot plot) and interquartile range of normalized kinetochore intensity values. n refers to individually measured kinetochores (left to right: 1032, 1171, 640,

1142). The experiment was performed three times. (**F**) Least-square linear fitting of data points for each kinetochore of the Zwilch intensity on the y-axis and the EGFP intensity on the x-axis of the experiment shown in (D). (**G**) Quantification of MAD1 levels at kinetochores of the experiment shown in (D). Red bars represent median (value shown above scatter dot plot) and interquartile range of normalized kinetochore intensity values. n refers to individually measured kinetochores (left to right: 833, 913, 749, 987). The experiment was performed three times. (**H-I)** Quantification of MAD1 and BUBR1 levels at kinetochores of the experiment shown in **Fig. 3H**. Red bars represent median (value shown above scatter dot plot) and interquartile range of normalized kinetochore intensity values. n refers to individually measured kinetochores (H left to right: 1882, 1375, 345, 1408, 1353, 2080 I left to right: 467, 399, 209, 181, 385, 413). The experiment was performed three times. (**J**) Quantification of Zwilch levels at kinetochores of stable DLD-1 cells treated as indicated in **Fig. 3G**. Red bars represent median (value shown above scatter dot plot) and interquartile range of normalized kinetochore intensity values. n refers to individually measured kinetochores (left to right: 679, 512, 1343, 963). The experiment was performed three times.

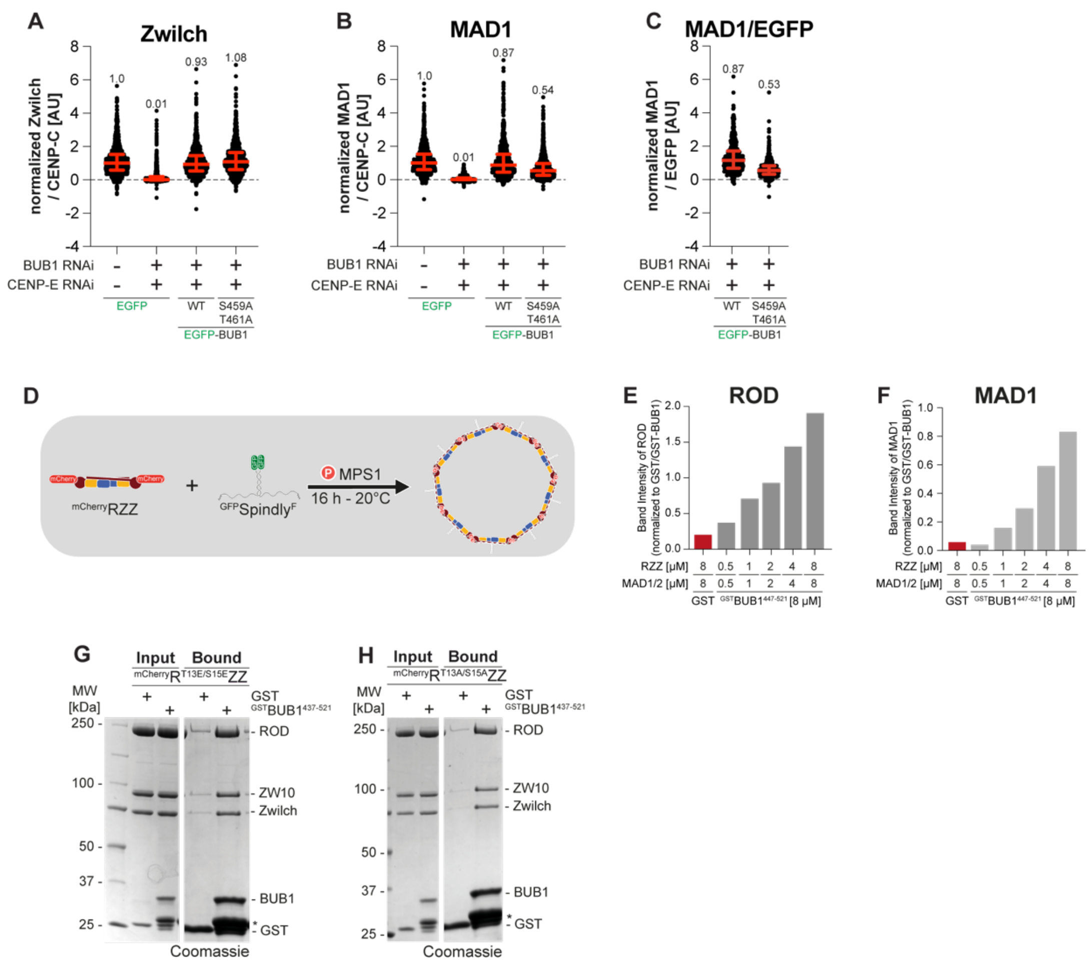


**Fig. S7.** (**A-B**) Quantification of Zwilch and MAD1 levels at kinetochores of stable DLD-1 cells treated as indicated in **Fig. 3G**. Red bars represent median (value shown above scatter dot plot) and interquartile range of normalized kinetochore intensity values. n refers to individually measured kinetochores (A left to right: 1445, 1059, 1028, 1012 B left to right: 1424, 786, 944, 1145). The experiment was performed two times. (**C**) Ratio of the MAD1/EGFP levels at kinetochores of the experiment quantified in (A-B). Red bars represent median (value shown above scatter dot plot) and interquartile range of normalized kinetochore intensity values. n refers to individually measured kinetochores (left to right: 944, 1145). (**D**) Schematic of RZZS ring formation used in the experiment depicted **Fig. 5C**. (**E-F**) Quantification of the ROD/MAD1 band intensity normalized to the bait signal from the SDS-PAGE depicted in **Fig. 5B**. (**G-H**) SDS-PAGE analysis of a pulldown assay with either GST or GST-tagged BUB1$^{437\text{-}521}$ as bait, and mCherry-RZZ constructs as prey. The experiment was performed two times.

**Table S1.**
List of the siRNA oligos used in this study.

| Target mRNA | Sequence (3'-5') | Concentration | Source |
|---|---|---|---|
| BUB1 | Dharmacon: GGUUGCCAACACAAGUUCU | 50 nM | *Krenn et al., 2014* |
| CENP-E | Dharmacon: AAGGCUACAAUGGUACUAUAU | 60 nM | *Ciossani et al., 2018* |
| KNL1 | Invitrogen:<br>HSS183683: CACCCAGUGUCAUACAGCCAAUAUU<br>HSS125942: UCUACUGUGGUGGAGUUCUUGAUAA<br>HSS125943: CCCUCUGGAGGAAUGGUCUAAUAAU | 60 nM | *Krenn et al., 2014* |
| Ndc80C | Sigma-Aldrich:<br>siHEC1: GAGUAGAACUAGAAUGUGA<br>siSPC24: GGACACGACAGUCACAAUC<br>siSPC25: CUACAAGGAUUCCAUCAAA | 20 nM | *Kim & Yu, 2015* |

**Table S2.**
List of the primary antibodies used in this study.

| Epitope | Species | Dilution | Manufacturer (cat. number) |
|---|---|---|---|
| BUB1 | Rabbit polyclonal | 1:500 | Abcam, #ab9000 |
| BUBR1 | Rabbit polyclonal | 1:1000 | Thermo Scientific #720297 |
| CENP-C | Guinea pig polyclonal | 1:1000 | MBL, #PD030 |
| CENP-E | Rabbit monoclonal | 1:200 | Abcam, #ab133583 |
| HEC1 | Mouse monoclonal | 1:1000 | Abcam, #ab3613 |
| KNL1 | Rabbit polyclonal | 1:750 | made in-house, #SI0787 |
| MAD1- DyLight488 | Mouse monoclonal | 1:200 | made in-house, Clone BB3-8 |
| MAD1-DyLight550 | Mouse monoclonal | 1:200 | made in-house, Clone BB3-8 |
| Zwilch | Rabbit polyclonal | 1:750 | made in-house, #SI520 |

**Table S3.**

List of the secondary antibodies used in this study.

| Name | Species | Manufacturer (cat. number) |
|---|---|---|
| α-mouse Alexa Fluor 488 | Goat | Invitrogen A11001 |
| α-mouse Rhodamine Red | Goat | Jackson Immuno Research 115-295-003 |
| α-rabbit Alexa Fluor 488 | Donkey | Invitrogen A21206 |
| α-rabbit Rhodamine Red | Donkey | Jackson Immuno Research 711-295-152 |
| α-guinea pig Alexa Fluor 647 | Goat | Invitrogen A-21450 |
